# Genome-wide nascent transcription profiling uncovers gene regulatory elements and dynamic immune responses in the major arbovirus vector *Aedes aegypti*

**DOI:** 10.1101/2023.11.03.565578

**Authors:** Femke A.H. van Hout, Samu V. Himanen, Anniina Vihervaara, Pascal Miesen

**Affiliations:** Department of Medical Microbiology, Radboud University Medical Center, Nijmegen, The Netherlands; Department of Gene Technology, Science for Life Laboratory, KTH Royal Institute of Technology, Stockholm, Sweden

**Keywords:** *Aedes aegypti*, enhancer, innate immunity, *in vivo* PRO-seq, midgut, microbiome, nascent RNA profiling, promoter, PRO-seq, transcription dynamics

## Abstract

*Aedes aegypti* mosquitoes are the principal vectors for epidemic arboviruses, including dengue, Zika, and chikungunya viruses. In these insect vectors, the concerted action of immune pathways shapes virus replication and transmission; yet, mechanistic and dynamic understanding of the transcriptional immune responses in mosquitoes remain limited. To address this knowledge gap, we profiled nascent transcription in mosquito cells and midgut tissue using Precision Run-On sequencing (PRO-seq). We identified exact transcription start nucleotides (TSNs), characterized promoter architectures, and generated the first genome-wide list of putative enhancers in *Aedes aegypti*, substantially expanding the regulatory annotation of this non-model organism. To investigate dynamic immune responses, we stimulated Aag2 cells with heat-inactivated *E. coli* and measured RNA synthesis and mRNA levels from matched samples. Bacterial stimulation induced rapid and temporally organized transcriptional responses associated with distinct biological functions and transcription factor motifs. Integration with mRNA-seq revealed that temporal patterns of mRNA accumulation were shaped by RNA synthesis, gene length, and stability, indicating multilayered regulation of mosquito immune responses. Implementing PRO-seq for mosquito midgut enabled characterization of tissue-specific transcription and TSN usage. Importantly, we demonstrate for the first time that PRO-seq captures *in vivo* microbial transcription within an organism, co-profiling host and microbiome nascent RNA synthesis. Altogether, this study substantially expands the regulatory annotation and functional genomic landscape of *Aedes aegypti*, reveals multilayered regulation of mosquito immune responses, and extends transcriptional profiling to host and microbiome activity *in vivo*. These insights advance understanding of gene regulation and immunity in this important arbovirus vector.

## Background

Emerging and re-emerging epidemic arthropod-borne (arbo) viruses, including dengue, Zika, and chikungunya viruses, pose significant threats to global health with nearly half of the world’s population currently at risk of infection^1,2^. The main vector for arbovirus transmission is *Aedes (Ae.) aegypti*, an urban-adapted mosquito that originated in Africa, but due to global trade and travel is now endemic in tropical and subtropical regions all over the world^5^. The ability of mosquitoes to transmit viruses, termed vector competence, depends on successful infection and replication of the virus within the insect host^6–8^. Consequently, mosquito-intrinsic pathways, including stress responses and immune defenses, determine virus replication and transmission efficiency into humans^9,10^. In addition to transmitting arboviruses, mosquitoes harbor diverse environmental microbes, including bacteria, fungi, and insect-specific viruses. This results in an intriguing triangular interaction between the mosquito, its microbiota, and human-pathogenic viruses, in which microbial communities can directly or indirectly influence virus infection and transmission^11–14^. It is therefore crucial to obtain a holistic view of mosquito host responses to various microbes.

Microbe recognition triggers host signaling pathways that coordinate gene expression and antimicrobial responses. In mosquitoes, these include the nuclear factor kappa B (NFκB)-related *Toll* and *Immune deficiency* (Imd) pathways, as well as the *Janus kinase-Signal Transducers and Activators of Transcription* (JAK-STAT) pathway^15,16^. Additionally, insulin signaling, c-Jun N-terminal kinase (JNK), and Heat Shock Factor (HSF1) have been implicated in antimicrobial defense reponses^17–19^. However, mechanistic understanding of transcriptional regulation during mosquito immune responses remains limited. Transcriptional responses are highly dynamic processes that involve rapid and coordinated changes in RNA synthesis. Conventional mRNA-sequencing (mRNA-seq) methods report steady-state levels of polyA-enriched transcripts, which are determined not only by transcriptional activity but also by post-transcriptional processes, including transcript stability and microRNA (miRNA)-mediated silencing. In contrast, nascent RNA methods directly capture ongoing RNA synthesis and measure dynamic changes in gene expression at high sensitivity and resolution^20^. Integration of these complementary approaches can therefore provide insight into multiple layers of gene regulation, linking transcriptional induction and repression to post-transcriptional regulation and mRNA expression levels.

Among nascent RNA approaches, Precision Run-On sequencing (PRO-seq) enables genome-wide and strand-specific mapping of actively engaged RNA polymerases at nucleotide resolution^21^. PRO-seq is based on a run-on reaction, during which transcriptionally engaged RNA Polymerases incorporate a single labeled ribonucleotide into the 3’-end of growing RNA chains, labeling the active sites of transcription. Consequently, PRO-seq has provided critical insights into the mechanisms that control RNA polymerase II (Pol II) at genes and enhancers and allows tracking Pol II progression from the precise initiating nucleotide through the rate-limiting steps of transcription^20^.

This high-resolution view on engaged Pol II across the genome has been particularly valuable for disentangling the dynamic transcriptional responses to diverse stimuli, including heat shock^22^, oxidative stress^23^, cytokines^24^, and pathogen challenge^25^. Moreover, PRO-seq captures transcription initiation and enhancer-associated transcription, enabling high-resolution mapping of transcription start nucleotides (TSNs, +1 nt) and enhancer RNAs (eRNAs)^26–28^. Such information is particularly valuable for non-model organisms such as *Ae. aegypti*, whose regulatory landscape remains incompletely characterized.

In this study, we dissect transcriptional regulation in *Ae. aegypti*, a major vector of arboviruses. By generating the first genome-wide maps of ongoing transcription, we identify exact transcription start nucleotides and provide a catalogue of putative enhancers in this species. Following bacterial stimulation, we identify temporally organized waves of transcriptional responses that associated with distinct transcription factor motifs and biological functions, including rapid AP-1 and NF-kB-associated responses followed by a secondary ER stress-related response. Integration with matched mRNA-seq further uncovered that gene activation, transcript production time, and RNA stability contribute to the temporal organization of immune gene expression programs. Finally, profiling of mosquito midguts captures host and microbiome RNA synthesis *in vivo*, providing insight into microbial activity within the mosquito gut environment. We recover transcription signatures simultaneously from host and its microbiome, including taxa such as *Elizabethkingia*, previously associated with *Ae. aegypti* midguts. Altogether, these findings expand the functional genomic landscape of *Ae. aegypti*, demonstrate PRO-seq to capture microbiome and host nascent transcriptomics, and creates new avenues for uncovering mechanisms of mosquito gene regulation, immunity, and vector competence, with broad implications for understanding and controlling arbovirus transmission.

## Results

### Profiling transcription start nucleotides and core promoter architecture in *Ae. aegypti*

PRO-seq data provides a powerful resource for annotating regulatory elements that govern transcription and is particularly valuable for non-model organisms where genome annotations remain incomplete. To study transcriptional regulation in *Ae. aegypti,* we optimized PRO-seq for the Aag2 cell line (Fig. S1A), and generated libraries from naïve cells and cells exposed to heat-killed bacteria. Prior to analyzing the effects of immune challenge, we harnessed PRO-seq to map the exact transcription start nucleotides (+1 nts; TSNs) on a genome-wide scale^26^ (Fig. 1A-C). This analysis identified 12,881 TSNs (Fig. S2A, S2B, Supplementary Table 1), of which only a remarkably small fraction of less than1% coincided with annotated transcription start sites (TSSs) (Fig. 1B, Fig. S2C), highlighting the lack of nucleotide resolution in the current annotation of gene starts. The PRO-seq-defined TSNs showed an average downstream shift of +55 nt (Fig. 1C, D), a pattern also reported in human and dog cells^26,27^. Since such offsets in annotated start positions could obscure cis-regulatory elements, we evaluated core promoter motifs relative to either the annotated TSSs or PRO-seq-defined TSNs. Specifically, we assessed the positioning of the TATA box, the Initiator (Inr) motif, and the downstream promoter element (DPE) (Fig. 1E, upper panel). TATA boxes were detected in approximately 300 promoters for both the current annotation TSSs and the PRO-seq-defined TSNs. However, enrichment at the expected position ∼30 nt upstream of the initiation site^3^ was observed only when aligned to PRO-seq-defined TSNs (Fig. 1E, left). Inr and DPE motifs were more frequently identified when using PRO-seq-defined TSNs, with their occurrence increasing by 39% and 29%, respectively (Fig. 1E). Strikingly, while neither motif showed a clear positional enrichment relative to current TSS annotations, alignment relative to PRO-seq-defined TSNs revealed positioning of these core promoter elements with nucleotide precision (Fig. 1E, middle and right). The central adenine of the Inr motif peaked exactly at the +1 nt position and the DPE was located at +28 nt relative to the TSN (Fig. 1E, middle and right; Fig. 1F), consistent with its positioning described in other systems^3^. These results demonstrate that TSN calling by PRO-seq substantially improves the existing *Ae. aegypti* genome annotations, providing a high-resolution map of transcription initiation and promoter architecture.

**Figure 1.**
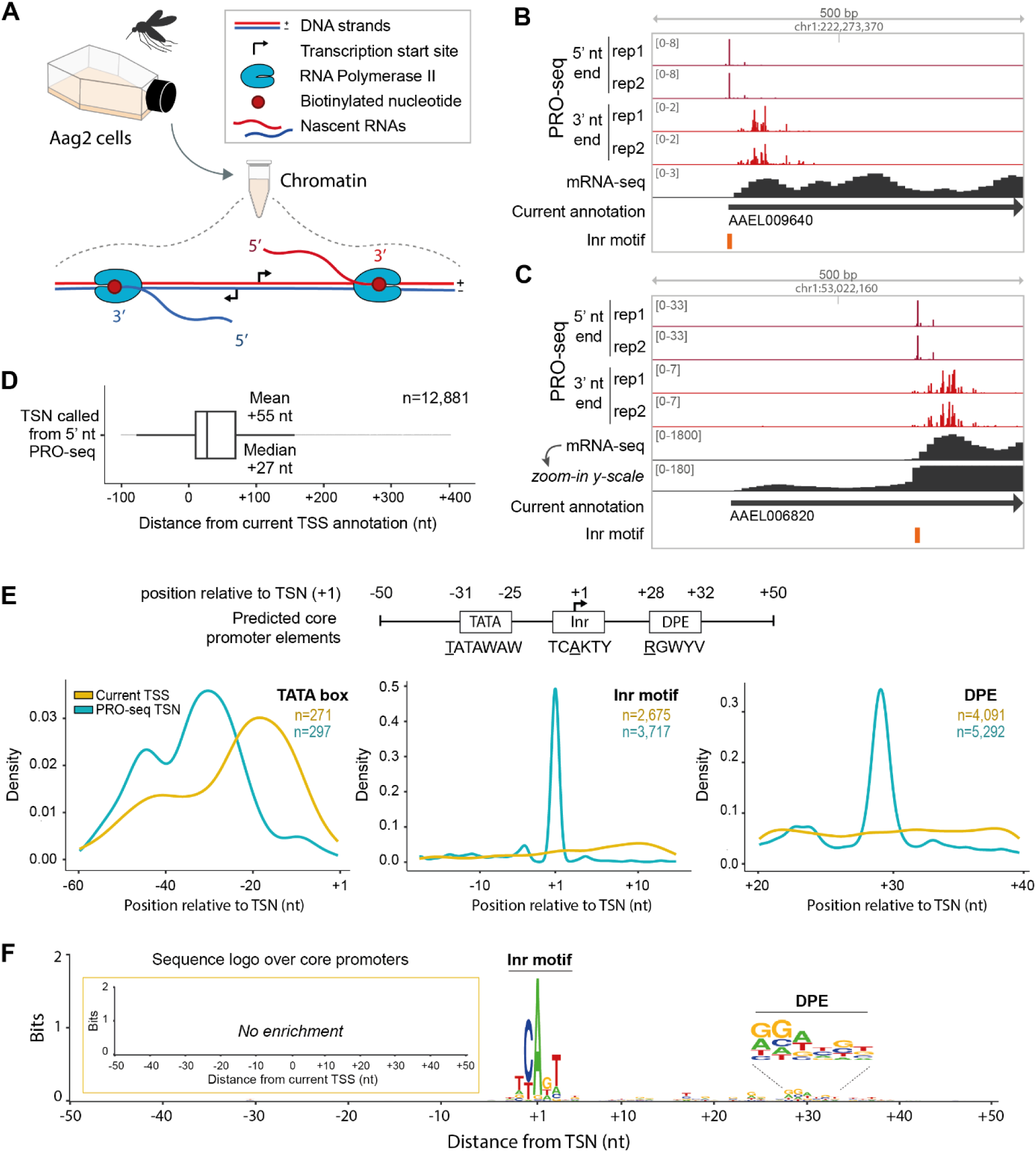
Genome-wide annotation of transcription start nucleotides (+1 nt, TSNs) and core promotor elements in *Ae. aegypti*. **A)** Schematic representation of engaged Pol II complexes and +1 nts identified with PRO-seq. Biotinylated ribonucleotides are incorporated into the 3’-ends of nascent RNAs by Pol II, enabling the purification of nascent transcripts. The 3’ ends of nascent RNAs represent the genomic positions of engaged Pol II, while the 5’ ends enrich at the precise transcription start nucleotide. **B+C)** Genome browser tracks of PRO-seq and mRNA-seq around start of genes. An example for PRO-seq 5’-nt coverage coinciding with (B) and localizing downstream of (C) the currently annotated transcription start sites are shown. The position of the initiator motif is indicated with an orange bar. **D)** Distance between the transcription start nucleotide based on 5’-nt PRO-seq coverage (TSN) and current start site annotation (TSS). **E)** Upper panel: Core promoter elements and their positions around the transcription start as described for *D. melanogaster*^3^, including TATA box, Initiator (Inr) motif and downstream promoter element (DPE). Lower panels: Density distribution of core promoter elements around transcription start sites based on either the current gene annotation (yellow) or PRO-seq data (blue). The position of the underlined nucleotide in the sequence motifs is depicted. *n* indicates the number of identified core promotor elements in each dataset. **F)** Sequence motif generated for a 100 nt window around TSNs called by 5’-nt PRO-seq peaks (large panel) and the current TSS gene annotation (insert).

### Genome-wide analysis of engaged RNA Pol II reveals promoter-proximal pausing and largely unidirectional transcription at *Ae. aegypti* promoters

To map the positions of transcriptionally engaged Pol II across the *Ae. aegypti* genome, we analyzed the distribution of 3’-nt ends of PRO-seq reads (Fig. 2A, Fig. S3A). At active genes, PRO-seq profiles typically reveal promoter-proximal pausing, productive elongation on gene bodies, slow-down of transcription at cleavage and polyadenylation site (CPS), and subsequent termination of transcription (Fig 2A). Additionally, PRO-seq profiles detect divergent transcription from promoters and enhancers, which is particularly prominent in mammalian cells (Fig. 2A)^29^. In *Ae. aegypti,* the majority of PRO-seq reads mapped to annotated genes (Fig. 2B, upper panel). Approximately 20% of the reads localized to intergenic transposable elements (Fig. 2B, upper panel), reflecting the high content of actively transcribed transposons in the *Ae. aegypti* genome^30^. About one quarter of the engaged Pol II could not be assigned to currently annotated genomic features. Notably, this fraction decreased to ∼5% when reads mapping to the opposite strand were included (Fig. 2B, upper panel), indicating antisense transcription at genes and transposable elements. The remaining unassigned signal likely reflects transcription from currently unannotated functional regions, such as enhancers, which are not yet included in the *Ae. aegypti* genome annotation.

**Figure 2.**
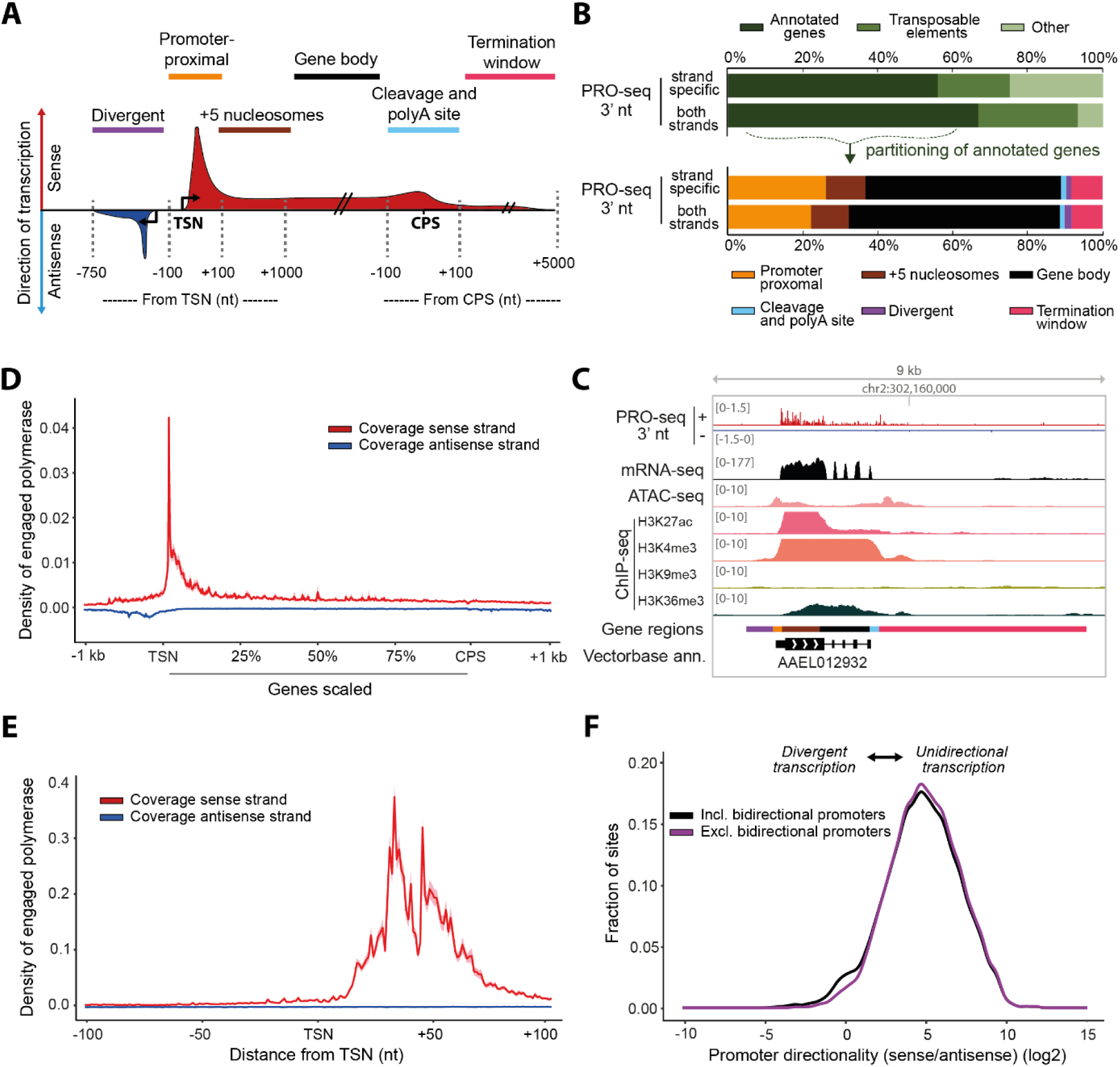
Genome-wide distribution of engaged RNA Pol II in *Ae. aegypti*. **A)** Schematic illustration of PRO-seq signal derived from 3′-ends of reads (PRO-seq 3′-nt) across genes. The subdivision into six functional gene regions used for the analysis in B is shown. TSN indicates transcription start nucleotide and CPS indicates cleavage and polyadenylation site. **B)** Upper panel: Percentage of PRO-seq 3’-nt coverage (engaged Pol II) mapping to genes, transposable elements (TE) and other regions. The analysis was either restricted to reads mapping in the same orientation as the gene/TE (strand-specific) or also included reads mapped to the opposite strand of the feature (both strands). Lower panel: Relative distribution of PRO-seq 3’-nt coverage across genes subdivided into the six different functional regions (Fig. 2A). **C)** Genome browser views of PRO-seq 3’-nt, mRNA-seq, ATAC-seq and ChIP-seq data for representative transcriptionally active gene, with gene regions as indicated in A. **D)** Metaplot of PRO-seq 3’-nt coverage over all annotated genes (scaled, bin size of 10 nt), including 1 kb unscaled region on either end. **E)** Metaplot of PRO-seq 3’-nt coverage over the 200-nt window around TSNs (non-scaled, bin size of 1 nt). **F)** Promoter directionality shown as log2-transformed ratio of sense to antisense reads around transcription initiation sites, calculated as in^4^.

To characterize Pol II progression through genes, we partitioned genes into six functional regions reflecting key transcriptional stages (Fig 2A, C, Fig. S3A)^29^. This analysis showed that most of the engaged Pol II in Aag2 cells was distributed along gene bodies (Fig. 2B, lower panel), including the early coding region around the first nucleosomes where Pol II proceeds slow and recruits elongation factors^31,32^. In *Ae. aegypti*, Pol II accumulation was prominent downstream of TSNs, consistent with promoter-proximal Pol II pausing (Fig. 2B, lower panel; Fig. 2D), a key regulatory step that controls the release of Pol II into productive elongation^20,31,33–35^. More specifically, we found two Pol II pause peaks around +30 nt and +50 nt (Fig. 2E), consistent with the early and late pause sites detected in metazoan species^26,36^.

While promoter-proximal pausing was wide-spread in *Ae. aegypti*, antisense transcription upstream of promoters was generally limited (Fig. 2D), indicating that transcription at *Ae. aegypti* promoters is largely unidirectional rather than divergent. To quantify promoter directionality, we determined the ratio of sense over antisense PRO-seq signal around the TSNs^4^ (Fig. 2F). The vast majority of promoters showed high directionality, with 20-30 fold higher sense than antisense transcription, indicating predominantly unidirectional promoter architecture in *Ae. aegypti.* This resembles promoter architecture described in *Drosophila* but is in contrast to the widely observed divergent transcription at mammalian promoters^4,27,37–40^.

Nevertheless, the directionality distribution showed a small shoulder around zero, indicating a minor subset of promoters with comparable levels of sense and antisense transcription (Fig. 2F). Inspection of these loci revealed two distinct configurations: divergent transcription from promoters driving a single gene (Fig. S3B), and transcription from bidirectional promoters, where two divergently oriented genes start from the same promoter region (Fig. S3C). Exclusion of these bidirectional gene pairs strongly reduced the shoulder around zero (Fig. 2F, purple line), indicating that bidirectional promoter organization accounts for a substantial fraction of the observed antisense signal. However, individual examples of divergent transcription at single-gene promoters remained (Fig. S3B), suggesting that rare cases of divergent transcription occur in *Ae aegypti*. These divergent promoters displayed distinct patterns of epigenetic marks than the bidirectional promoters of gene pairs (Fig. S3B-C). In addition to unidirectional transcription at the start of the genes, also the gene ends in *Ae. aegypti* showed distinct profiles to mammalian cell. In *Ae. aegypti*, we detected minimal Pol II accumulation near cleavage and polyadenylation sites (CPS) (Fig. 2D), suggesting that transcription termination in mosquito cells is less associated with slowing down of Pol II.

Together, these data establish key features of transcription in *Ae. aegypti*, showing highly positioned core motifs (Fig. 1), largely unidirectional gene transcription, pronounced promoter-proximal pausing at early and late pause regions, and minimal Pol II accumulation at gene ends (Fig. 2). In addition, transcription outside annotated genes highlights widespread transcription of transposable elements and suggests the presence of uncharacterized regulatory elements such as enhancers.

### Nascent transcription reveals putative enhancers and enhancer clusters in *Ae. aegypti*

Beyond promoters, transcription can also initiate at enhancers, producing short-lived enhancer RNAs (eRNAs)^39,41^ that correlate with enhancer activity (reviewed by Tippens, *et al.* (2018)^42^). By inspecting transcription outside of promotors, nascent RNA sequencing has been widely used to profile eRNAs across vertebrate and invertebrate organisms, providing an important contribution to the identification of enhancers^22,27,28,37,42–45^.

To map putative enhancer elements on a genome-wide scale, we identified transcriptional regulatory elements (TREs) *de novo* using dREG^44^. After excluding TREs overlapping promoters (Gene TSS +-500 nt), almost 35,000 enhancer candidates were identified. Through integration with ATAC-seq, indicating accessible chromatin, and ChIP-seq for H3K27ac, a histone mark used as a hallmark of active enhancers, we refined this set to 18,586 putative enhancers (Fig. 3A, S4A, Supplementary Table 2). For these, we identified the TSN, considering both genomic strands separately (Fig. S4B) and found, similar to promotors, strong enrichment for the Initiator (Inr) motif (Fig. 3B). We next compared transcriptional and chromatin features between gene promotors and putative enhancers. To enable direct comparison with promoters, enhancer regions were aligned according to their dominant direction of transcription (Supplementary Table 3). Enhancers exhibited overall lower levels of transcription and shorter nascent RNAs than promoters (Fig. 3C). Yet, antisense transcription was more prevalent at enhancers than promoters (Fig. 3C-D; Fig. S4C), consistent with prior studies in human, fruit fly, and dog^4,22,27,37^.

**Figure 3.**
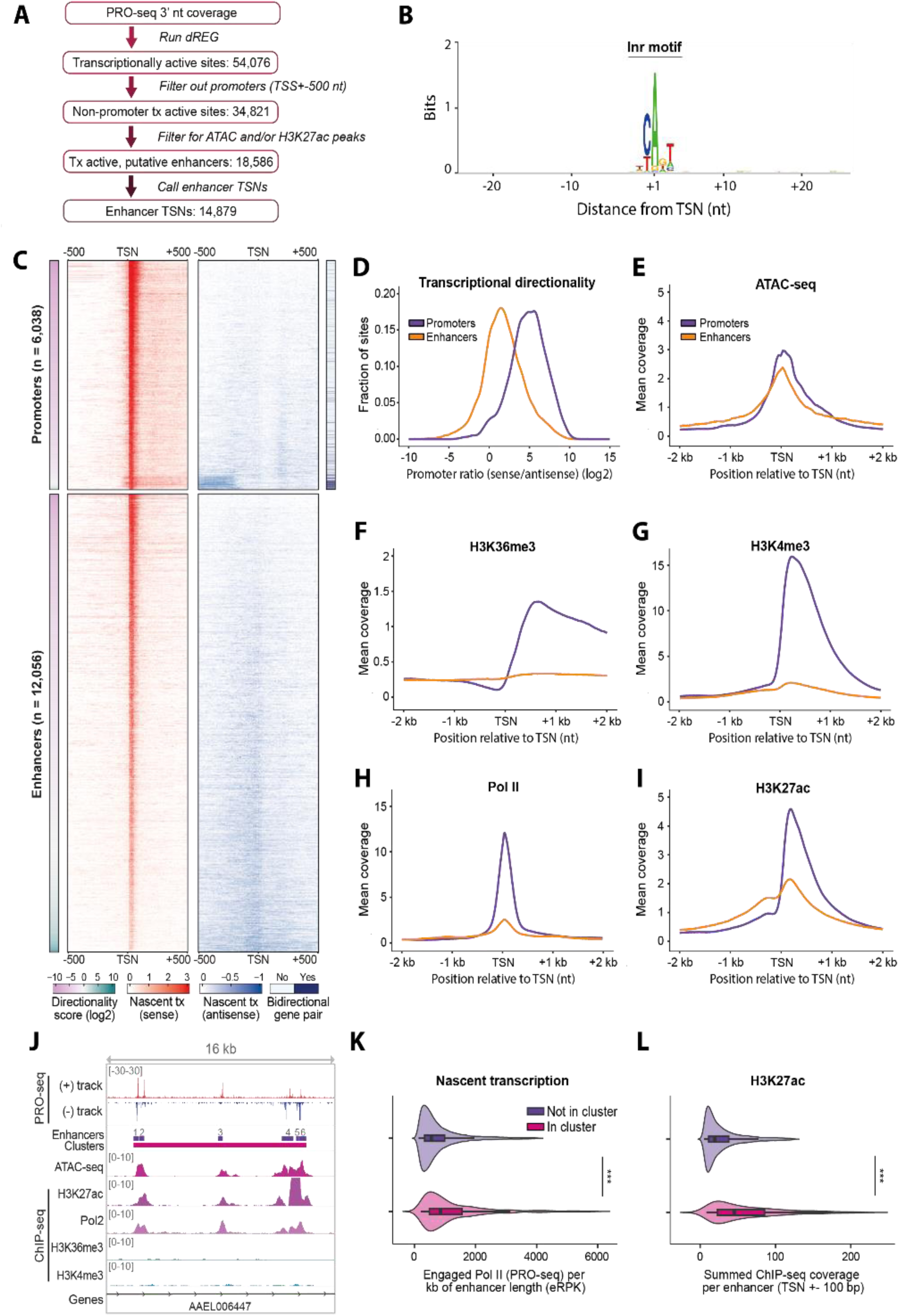
Transcription and chromatin state at enhancers in the *Ae. aegypti* genome. **A)** Strategy to identify enhancers in the *Ae. aegypti* genome. The dREG algorithm was used to identify all transcriptional regulatory elements (TREs). Non-promoter TREs were selected and further filtered for overlap with ATAC and/or H3K27ac peaks to generate the list of transcriptionally active, putative enhaners (Supplementary Table 2). For each putative enhancer, transcription start nucleotides (TSNs) were identified on both strands, and the dominant TSN was selected (see also Fig. S4B). **B)** Sequence motif generated within a 50-nt window around the TSNs called for putative enhancers. **C)** Heatmaps of nascent transcription (PRO-seq 3’-nt signal), shown as sense (red) and antisense (blue) signal. Both heatmaps show the regions in the same order, sorted based on transcriptional directionality. **D)** Distributions of transcriptional directionality (log2 ratio of sense to antisense reads) at promoters and enhancers, calculated as in Core *et al.* (2012)^4^. **E-I)** Metaplots of chromatin accessibility (E: ATAC-seq), Polymerase II ChIP-seq (H: Pol2), and histone marks (F: H3K36me3, G: H3K4me3, I: H3K27ac), around TSNs (+-500 bp) of enhancers and promoters. Corresponding heatmaps are displayed in Figure S4D. **J)** Genome browser view of an enhancer cluster with PRO-seq, ATAC-seq, and ChIP-seq profiles (Pol II, H3K27ac, H3K36me3, H3K4me3). Example region at chr2:437,690,890-437,703,000. All identified enhancer clusters are listed in Supplementary Table 4. **K+L)** Violin plots with embedded boxplots; values were trimmed to the 5^th^-95th percentile range for visualization; statistical significance was assessed on the full dataset using two-sided Wilcoxon rank-sum tests with FDR correction: ns, FDR ≥ 0.05; *FDR < 0.05; **FDR < 0.01; ***FDR < 0.001. **K)** Nascent transcription at enhancers within *versus* outside enhancer clusters, quantified as density of engaged Pol II per kilobase of enhancer length (eRPK). **L)** Summed H3K27ac coverage at enhancers within *versus* outside enhancer clusters, measured as total coverage over TSN +-100 bp windows.

Chromatin profiling revealed distinctive features as well as similarities between enhancers and promoters. Whereas they showed comparable chromatin accessibility (Fig. 3E, Fig. S4D), the classical gene-associated marks H3K36me3 and H3K4me3 were strongly enriched at promoters (Fig. 3F-G, Fig. S4D). Notably, however, H3K4me3 has been reported to occur at the most active enhancers in flies and mammals, scaling with the levels of Pol II rather than the region’s regulatory function^28^. Consistent with this, we observed H3K4me3 enrichment at the most highly transcribed mosquito enhancers (Fig. S4E). Furthermore, Pol II occupancy, determined through ChIP-seq, was generally higher at promoters than at enhancers, reflecting higher transcriptional activity. Enrichment of H3K27ac was prominent at both classes, but with notable differences in distribution (Fig. 3I, Fig. S4D). While promoter-associated H3K27ac signal was prominent downstream of the TSN, mirroring the strong directionality of gene transcription, enhancer-associated H3K27ac showed a more even distribution upstream and downstream of the TSN, in line with higher antisense transcription at enhancers.

To explore enhancer organization in the *Ae. aegypti* genome, we investigated the presence of enhancer clusters using eClusterer^27^. Clusters of closely spaced enhancers, also called super-enhancers, have been described in other species and are linked to coordinated regulation of transcription programs, particularly driving cell-type specific and cancer-associated genes^46,47^. We defined clusters as groups of ≥5 enhancers within a 12.5 kb window, extending the window if additional enhancers were found within a 2 kb distance^27^. We identified 124 clusters containing in total 735 enhancers (Fig. 3J, Supplementary Table 4). Consistent with previously reported properties of super-enhancers^27,46,47^, enhancers within clusters exhibited higher transcriptional activity (Fig. 3K) and higher levels of H3K27ac ChIP-seq, Pol II ChIP-seq, and ATAC-seq signal (Fig. 3L, S4F) than enhancers outside of clusters. Together, these analyses establish the first comprehensive annotation of putative enhancers and super-enhancers in *Ae. aegypti* cells, substantially expanding the landscape of gene regulatory elements in this key vector species.

### Transcriptome-wide estimates of RNA stability relate to miRNA targeting and gene function

Gene expression is shaped by both RNA synthesis and post-transcriptional processes. To investigate how these regulatory layers contribute to gene expression in *Ae. aegy*pti, we compared matched PRO-seq and mRNA-seq profiles generated from the same Aag2 samples. Whereas PRO-seq captures ongoing RNA synthesis, mRNA-seq quantifies steady-state levels of mature, polyadenylated transcripts.

Transcriptome-wide comparison revealed an overall correlation between RNA synthesis and mRNA abundance (Fig. 4A), indicating that RNA synthesis broadly predicts RNA expression levels. However, notable differences were observed between transcript classes. Highly transcribed RNAs with low mRNA levels are expected to constitute of unstable or non-polyadenylated RNAs. Indeed, histone mRNAs that lack poly(A) tails were predominantly captured by PRO-seq (Fig. S5A). Similarly, non-coding RNAs showed overall reduced mRNA-seq signal compared to coding transcripts (Fig. 4A), which may partially be explained by many ncRNAs lacking poly(A) tails. Yet, they also had overall lower production levels as measured by PRO-seq (Fig. 4A), in line with observations from other organisms^27,48^. Protein-coding transcripts detected by both methods showed a broad range of ratios between steady-state RNA levels (mRNA-seq) and nascent transcription (PRO-seq) (Fig. 4A, S5B). Previous studies have shown that this ratio provides transcriptome-wide estimates of RNA stability, with higher ratios reflecting increased transcript stability^48^. We therefore asked whether these inferred stability estimates reflected post-transcriptional regulation in mosquitoes and examined their relationship with miRNA targeting. miRNAs are conserved post-transcriptional regulators of gene expression that can modulate mRNA stability through sequence-specific targeting of transcripts. miRNA target sites can be classified into 6mer, 7mer-m8, and 8mer classes according to the extent and configuration of miRNA-target complementarity (see also Table 2), with 8mer sites generally mediating the strongest repression^49^.

**Figure 4.**
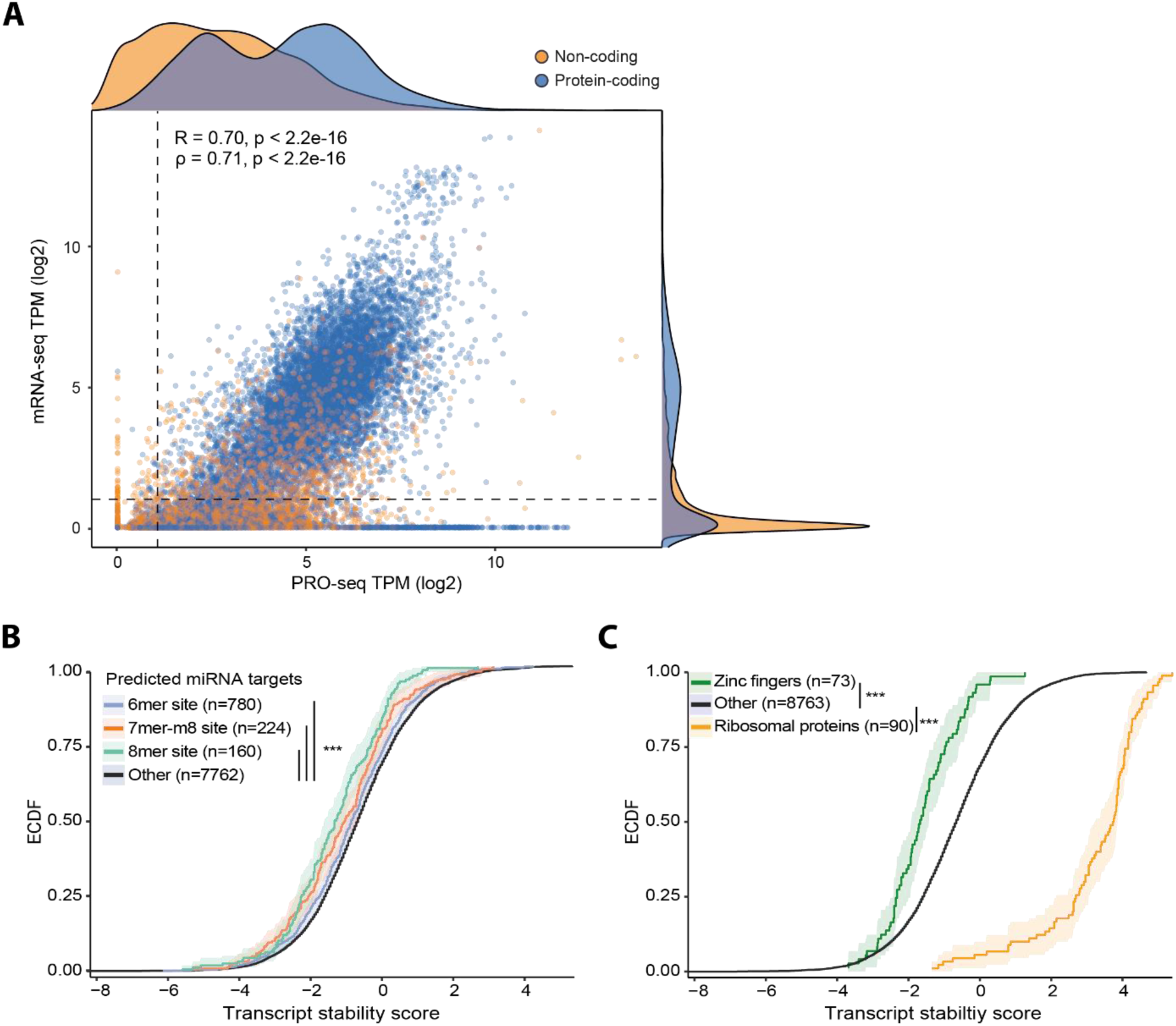
Estimating transcript stability from mRNA-seq and PRO-seq data reflects miRNA targeting and gene function. **A)** Scatter plot of mRNA-seq *versus* PRO-seq in Aag2 cells, displayed as log2 transcripts per million (TPM) per transcript (see Methods), with Pearson correlation coefficient (R) and Spearman’s rank correlation coefficient (ρ). Density distributions show PRO-seq coverage (top) and mRNA-seq coverage (right), split between protein-coding (blue) and non-coding (orange) transcripts. **B+C)** Empirical cumulative distribution functions (eCDFs) of estimated transcript stability for protein-coding transcripts with log2(TPM) > 1 in both assays (see also Fig. S5B). Shaded areas indicate 95% confidence intervals from 1,000 bootstrap replicates. **B)** Comparison of transcript stability for protein coding genes containing 3’ UTR target sites for the three most abundant miRNAs in Aag2 cells. Transcripts were grouped according to target site class (6mer, 7mer-m8, 8mer site) using transcripts without target site as control (for 6mer sites Δmedian = −0.16 and p = 6.3e-2, for 7mer-m8 sites Δmedian = −0.35 and p = 4.0e-3, for 8mer sites Δmedian = 0.6 and p = 4.3e-5, with p-values according to Kolmogorov-Smirnov test with Benjamini-Hochberg correction). **C)** Comparison of transcript stability for genes encoding zinc finger proteins or ribosomal proteins versus all other protein-coding genes (for zinc fingers Δmedian = −1 and p = 6.7e-8, for ribosomal proteins Δmedian = 4.41 and p = 8.2e-54, with p-values according to Kolmogorov-Smirnov test with Benjamini-Hochberg correction).

In *Aedes* mosquitoes, miRNAs have been implicated in regulation of diverse processes, including development, digestion, and responses to arbovirus infection^50^. Using previously published miRNA expression data from Aag2 cells^51^, we tested whether transcripts targeted by abundant miRNAs showed reduced stability. Transcripts containing predicted target sites for abundant miRNAs in their 3’UTR indeed had lower stability values, with progressively stronger effects observed for 6mer, 7mer-m8, and 8mer sites (Fig. 4B), consistent with the expected hierarchy of miRNA target-site efficacy. In contrast, transcripts containing target sites for lowly expressed miRNAs in Aag2 cells did not show reductions in stability (Fig. S5C), supporting the biological relevance of the stability estimates.

We next examined whether transcript stability was associated with gene function. Stable transcripts were enriched for housekeeping and homeostatic processes, whereas unstable transcripts were enriched for signaling pathways (Fig. S5D, E). In line with this, transcripts encoding ribosomal proteins displayed significantly higher stability estimates, whereas transcripts encoding zinc finger proteins showed lower stability values (Fig. 4C), consistent with findings in mammalian systems^48^. Altogether, these findings demonstrate that integration of mRNA-seq and PRO-seq data enables transcriptome-wide inference of RNA stability that reflects post-transcriptional regulation and gene function in mosquitoes.

### Transcriptional immune responses are temporally organized in *Ae. aegypti* cells

To investigate transcriptional reprogramming during mosquito antibacterial defense, we stimulated Aag2 cells with heat-inactivated *E. coli* and collected PRO-seq and mRNA-seq samples at 1 and 4 hours post stimulation (Fig. 5A). PRO-seq revealed a rapid transcriptional response that was more pronounced at 1 than 4 hours after immune stimulation (Fig. 5B, C, Supplementary Tables 5, 6). To investigate whether this response triggered distinct temporal programs, genes induced across the time course were clustered based on their transcriptional dynamics. We identified three major waves of induction: rapid but transient activation, rapid and sustained activation, and delayed activation (Fig. 5D). Functional annotation revealed biological segregation between the temporal clusters (Fig. 5E, Supplementary Tables 7-9). While the rapid but transiently induced genes were enriched for membrane trafficking and cytoskeletal-associated functions, the rapidly and sustained induced genes showed strong enrichment for innate immune genes, including antimicrobial peptides (Fig. 5E). In contrast, delayed-response genes were enriched for ER stress and protein folding pathways, suggesting the emergence of a secondary stress response following antimicrobial activation.

**Figure 5:**
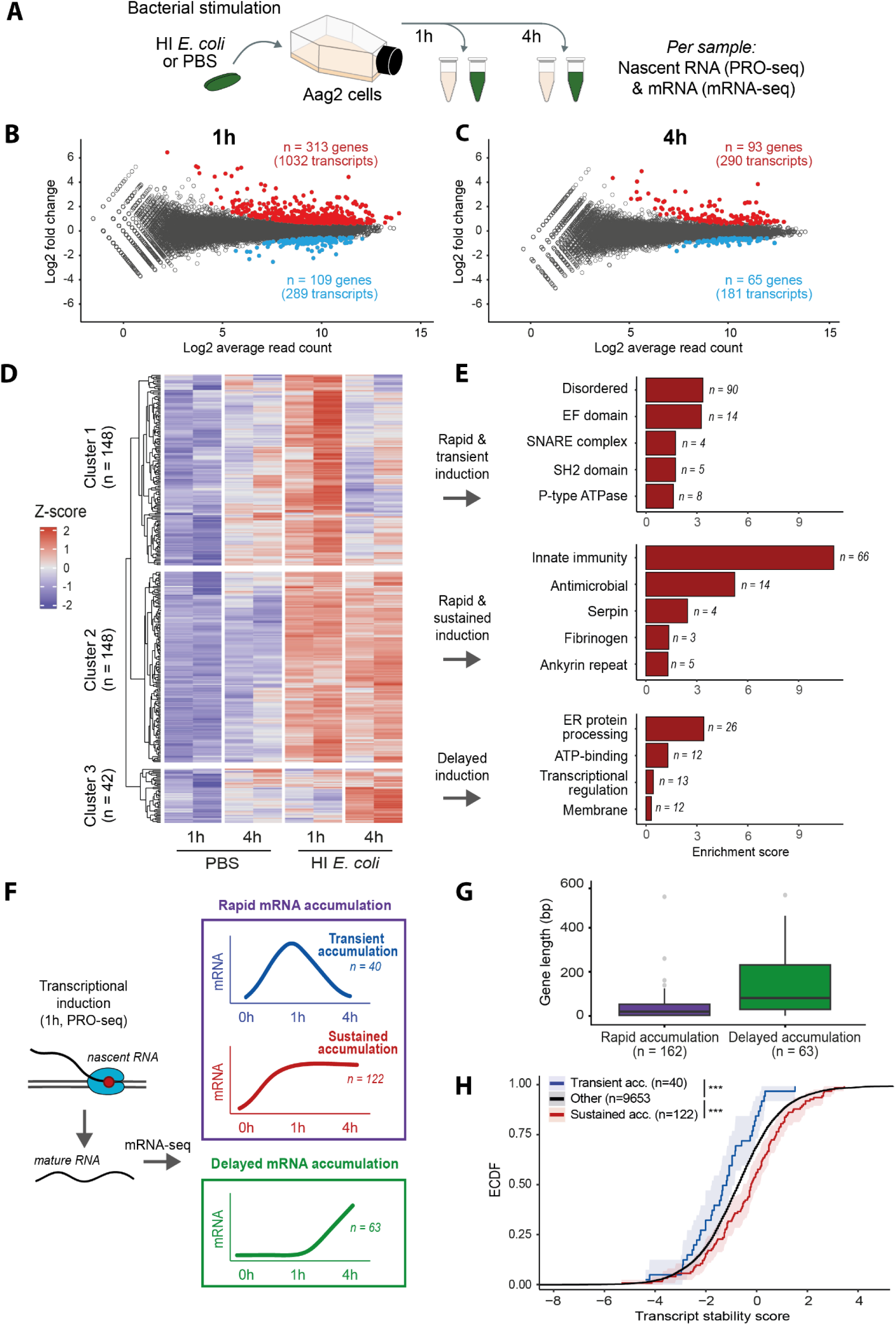
Temporal coordination of the Aag2 transcriptional response to immune stimulus. **A)** Experimental design: Aag2 cells were stimulated with heat-inactivated *E. coli,* followed by nascent RNA and mRNA collection after 1 and 4 hours for PRO-seq and mRNA-seq, respectively. **B+C)** MAplots of differentially transcribed genes at 1 hour (B) and 4 hours (C) after immune stimulus as measured by PRO-seq. **D)** Heatmap showing z-scored PRO-seq signal of genes transcriptionally induced at either 1 or 4 hours following stimulation. Genes were clustered using k-means clustering (k=3), with averages of biological duplicates as input. n indicates the number of genes in each cluster. **E)** Functional annotation of the temporal transcriptional clusters shown in (D) using DAVID. Top five enriched categories and the top-ranking term per category are shown; total enriched categories and terms are listed in Supplementary Tables 7-9. **F)** Schematic overview of mRNA accumulation groups defined for genes induced at 1 hour in PRO-seq. Genes with increased nascent RNA levels at 1 hour were first classified into rapid accumulation (increased mRNA levels at 1 hour) and delayed accumulation (increased mRNA levels only at 4 hours) groups. Rapid accumulation genes were further subdivided into transient (increased only at 1 hour) or sustained (increased at both 1 and 4 hours) mRNA accumulation groups. **G)** Gene length distribution of rapid and delayed mRNA accumulation groups defined in (F). **H)** Empirical cumulative distribution functions (eCDFs) of transcript stability for genes in the transient and sustained mRNA accumulation groups defined in (F). Shaded areas indicate 95% confidence intervals from 1,000 bootstrap replicates. Stability estimates were inferred from the ratio between PRO-seq and mRNA-seq signal from Fig. 4. For the transient group Δmedian = −0.516 and p = 1.22e-2 and for the sustained group Δmedian = 0.532 and p = 9.09e-4, with p-values according to Kolmogorov-Smirnov test with Benjamini-Hochberg correction.

To identify potential transcription factors driving these temporal waves, we performed motif enrichment analysis on promoter regions of each cluster. Promoters of the rapidly induced genes contained motifs for AP-1, a transcription factor complex controlled by MAPK pathways such as the JNK pathway. Instead, NF-κB motifs were associated only with the rapid and sustained gene cluster (Fig. S6A, Supplementary Tables 10-12). These findings demonstrate that even within swift responses and narrow timeframes, functional and organizational differences emerge from transcription factor-driven waves of gene expression. During the mosquito antibacterial response, these waves activate membrane and cytoskeletal genes, immune pathways, and stress responses.

Next, we examined how nascent transcriptional changes were reflected at the level of mature mRNAs (Fig. 5A). mRNA-seq revealed similar functional programs as PRO-seq but mRNA accumulation was generally more pronounced at 4 than at 1 hour (Fig. S6B-E, Supplementary Tables 13-17), likely reflecting the time delay between transcriptional changes and mRNA abundance, as determined by transcript production time, processing and turnover.

To identify features associated with distinct mRNA accumulation kinetics, all genes induced at 1 hour in PRO-seq were selected and classified according to their corresponding mRNA-seq profiles (Fig. 5F). While some transcriptionally activated genes had already accumulated mRNA by 1 hour, others showed increased mRNA levels only at 4 hours (Fig. 5F). Comparing these rapid versus delayed mRNA accumulation groups revealed a clear difference in gene length, reflecting the time Pol II needs to travel through the gene: rapidly accumulating transcripts were generally shorter than ∼120 kb, whereas transcripts with delayed mRNA accumulation were longer (Fig. 5G), consistent with estimated elongation rates of eukaryotic Pol II. These findings suggest that gene length contributes to the temporal ordering of immune gene expression programs.

Among genes with rapid mRNA accumulation, expression profiles further separated into transient and sustained mRNA expression groups (Fig. 5F). To determine whether transcript stability contributed to these distinct accumulation patterns, we used the stability estimates described in Fig. 4. Transiently expressed transcripts displayed lower stability, whereas sustained transcripts exhibited increased stability (Fig. 5H), indicating that mRNA stability acts alongside RNA synthesis to shape immune response timing. Altogether, these findings reveal that antibacterial gene expression programs are shaped not only by transcriptional induction, but also by gene length and stability, which together influence the timing and persistence of mRNA expression.

### PRO-seq on mosquito midgut tissues uncovers tissue-specific transcriptional activity

The mosquito midgut is a key barrier for arbovirus transmission, where immune responses and microbial interactions shape infection outcomes^6,13,52^. To characterize transcriptional regulation in this physiologically relevant tissue, we applied a PRO-seq protocol optimized for low-input material^53^ (Fig. 6A). The sequencing libraries, generated from approximately 500.000 midgut cells, captured transcriptional landscapes with clear promoter-proximal pausing and robust gene body signal (Fig. 6B). Importantly, transcription of known midgut-expressed genes was readily detected (Fig. 6C)^54^, confirming capture of tissue-specific transcriptional programs.

**Figure 6.**
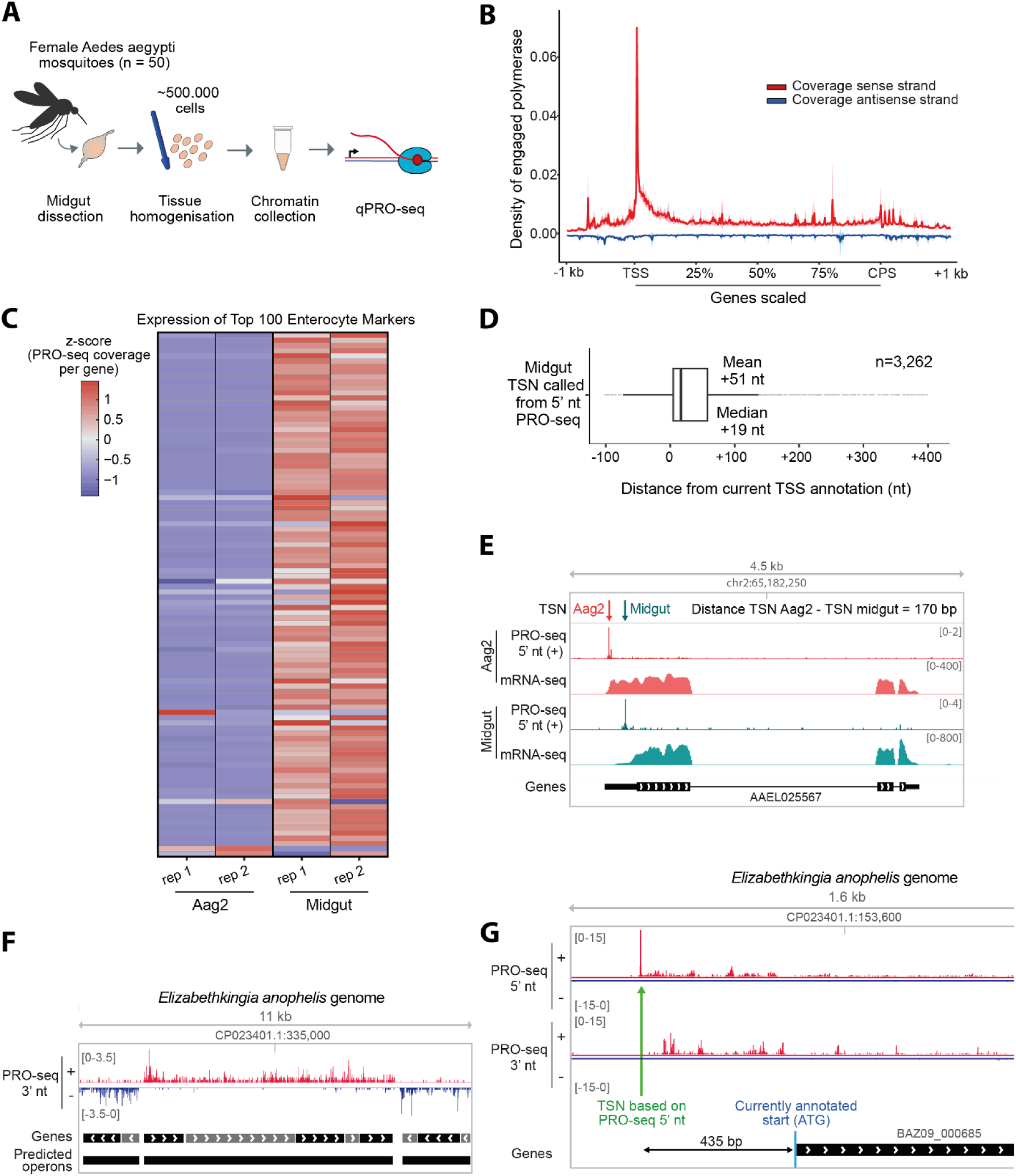
PRO-seq applied to mosquito midguts captures tissue-specific and microbial transcription. **A)** Experimental set-up: fifty female midguts (appr. 500.000 cells) were dissected and homogenized, after which chromatin was prepared as input for the qPRO-seq protocol. **B)** Metaplot of midgut PRO-seq 3’-nt coverage over all annotated genes (scaled, bin size of 10 nt), including 1 kb unscaled region on either end. **C)** Heatmap (z-scores) comparing expression in Aag2 *versus* midgut PRO-seq datasets, plotted for the top 100 enterocyte marker genes from the Mosquito Cell Atlas (see Methods). **D)** Distance between the transcription start nucleotide for midguts based on 5’-nt PRO-seq coverage and the current start site annotation for all transcripts passing the filtering steps as described in Fig. S7A. **E)** Genome browser view of PRO-seq 5′-nt and mRNA-seq tracks, illustrating differential TSN usage between Aag2 cells and midgut tissue. **F)** Genome browser view of PRO-seq 3’-nt coverage across *Elizabethkingia anophelis* genome. NCBI gene annotations (GCA_002023665.2) and operons predicted by Operon-mapper are shown. **G)** Genome browser view of *Elizabethkingia anophelis* PRO-seq 5′-nt and 3′-nt coverages together with the NCBI gene annotation (GCA_002023665.2). Current gene annotations lack 5’ UTR definitions and therefore start at the annotated translation start codon (ATG). Green arrow marks the PRO-seq 5′-nt peak upstream of the annotated start.

We next mapped transcription initiation in midgut tissue, pinpointing 3,262 TSNs with a mean deviation of 51 nt from current TSS annotations (Fig. 6D, S7A, Supplementary Table 18). To investigate whether transcription initiation differed between mosquito cells and midgut tissue, we compared midgut TSNs with those previously identified in Aag2 cells (Fig. 1). Among transcripts with a resolved TSN in midgut tissue, 20% (n = 643) lacked a corresponding TSN call in Aag2 cells, largely due to low expression in the cell line (Fig. S7B, D). Among transcripts with TSNs identified in both sample types, 73% initiated from identical nucleotides, whereas 27% exhibited differential TSN usage with a median shift of +53 nt (Fig. 6E, Fig. S7B-D). Such differences point to dynamic promoter selection across cellular contexts, which may alter responsiveness to distinct transcription factors and regulatory inputs. Resulting differences in transcript topology could affect RNA stability and translational output, potentially leading to differences in protein abundance or localization. For example, transcription from gene AAEL003193 initiated from a more downstream site in midguts than in Aag2 cells, resulting in a shorter midgut-specific isoform not yet annotated in the *Ae. aegypti* genome (Fig. S7E). Such a shorter isoform has been reported for the *Drosophila* ortholog (FBgn0016687, Nurf-38), indicating that PRO-seq identifies genuine examples of tissue-specific alterative TSN usage. Together, these findings show that *in vivo* transcriptional profiling captures tissue-specific transcriptional landscapes, insights which will be relevant for further studying mosquito-pathogen interactions in physiologically relevant tissues.

### *In vivo* PRO-seq co-profiles host and microbiome transcription

The mosquito midgut harbors a dynamic microbiome that can influence immune responses and pathogen susceptibility^55^. Because PRO-seq captures all actively engaged RNA polymerases within a sample without selection for its origin, we asked whether the midgut datasets also contained transcriptional signatures of resident microbiota. Reads not aligning to the mosquito genome were therefore taxonomically classified, revealing a diverse microbial community (Fig. S7F-H). Notably, these data demonstrate that PRO-seq simultaneously captures host and microbial transcription, enabling host-microbiome co-profiling in vivo.

Nascent profiling *in vivo* revealed a diverse microbial community (Fig. S7H), including bacterial genera previously reported for *Aedes* midguts, such as *Bacillus, Citrobacter, Escherichia* and *Klebsiella*^56–59^. While multiple genera were detected, the microbiome was dominated by *Elizabethkingia*, a genus previously reported to become highly abundant in mosquito midguts following blood feeding^57^. For the dominant bacterial species identified, *Elizabethkingia anophelis*, *in vivo* PRO-seq generated high-resolution coverage corresponding to transcriptional units (Fig. 6F), and indications of TSNs upstream of annotated coding sequences (Fig. 6G). This provides opportunities to refine promoter and operon architecture in densely packed microbial genomes, the annotation of which is currently often limited to the protein coding sequence, ignoring the non-coding regions with regulatory potential. By capturing host and microbiome transcription in parallel, the use of PRO-seq *in vivo* opens new avenues to investigate how transcriptional regulation and microbial activity in the midgut together influence mosquito infection outcomes and vector competence.

Taken together these findings illustrate how nascent RNA sequencing *in vivo*, captures the composition of complex biological systems, and profiles ongoing RNA synthesis across eukaryotic species and their prokaryotic symbionts.

## Discussion

Transcriptional regulation enables mosquitoes to coordinate immune responses to microbes, thereby influencing their capacity to transmit pathogens. In this study, we adapted PRO-seq to *Ae. aegypti* Aag2 cells and midgut tissue to map engaged Pol II at nucleotide resolution. By integrating nascent transcription profiling with chromatin and mRNA-seq datasets, we generated a comprehensive view of transcriptional regulation in this important disease vector. Our analyses refined genome annotation through identification of exact transcription start nucleotides (TSNs, +1 nts) and putative enhancers, revealed key features of mosquito transcriptional architecture, including unidirectional Pol II orientation at promoters and divergent transcription at enhancers. Additionally, we uncovered temporally organized antibacterial response programs, and simultaneously profile host and microbiome transcription programs *in vivo*. Together, these data provide a comprehensive framework to study gene regulation and host-microbe interactions in mosquitoes.

### Transcriptional architecture of mosquito genome at nucleotide-resolution

PRO-seq provided a detailed view of transcriptional architecture in *Ae. aegypti*, advancing the current genome annotation in this important disease vector. Mapping of +1 nts from the nascent RNAs revealed that current annotations often lack nucleotide precision, thereby obscuring positioning of core promoter motifs. Such a high resolution is critical for defining and interpreting the positioning of transcription factor binding sites, Pol II pausing, and chromatin organization, which remained mostly unexplored in this non-model organism. We found transcription at genes to be predominantly unidirectional and associated with strong promoter-proximal pausing. Similar architecture has been described in *Drosophila* promoters^4^, which contrasts the widespread divergent transcription seen at vertebrate promoters^27,38,60^. Divergent transcription has been associated with promoter composition, such as specific promoter elements, insulators, and chromatin features, such as CpG methylation^4,60–62^. Whether the unidirectional transcription observed in *Ae. aegypti* is linked to specific promoter elements, or relates to the limited DNA methylation present in dipteran insects^63^, remains to be investigated.

Beyond promoters, PRO-seq detected nascent transcription at putative enhancers, which lacked H3K36me3 and showed overall higher antisense transcription than promoters. These patterns are consistent with enhancer-associated transcription observed in other metazoans, strongly suggesting that our approach is able to identify functional enhancer elements^4,64^. Eventually, experimental validation of these putative enhancers will be important for understanding if and how these loci shape mosquito gene expression, immunity, and vector competence. Overall, these findings demonstrate how nascent transcription profiling can expand the functional genomic annotation for non-model species and open new opportunities to investigate gene expression regulation in mosquitoes.

### *Aedes aegypti* triggers dynamic transcriptional responses upon microbial challenge

Beyond defining transcriptional architecture, PRO-seq enabled direct quantification of antibacterial responses organized into temporally distinct transcriptional waves that were associated with different biological functions and transcription factor motifs. Rapid but only transiently transcribed genes were enriched for membrane trafficking and cytoskeletal functions, whereas the sustained genes associated with NF-κB-related motifs and antimicrobial effectors. In contrast, late induced genes were enriched for ER stress and protein-folding pathways, suggesting a secondary cellular stress response to support antimicrobial protein production and protein homeostasis. Similar temporal organization of immune and stress responses has been described in *Drosophila*^65^, indicating that transcriptional waves may represent a conserved feature of insect antibacterial immunity. Importantly, integration of PRO-seq with matched mRNA-seq demonstrated that mRNA accumulation is shaped by multiple regulatory layers, namely activation timing, transcript production time and RNA stability. Delayed mRNA accumulation was linked to longer gene lengths, whereas transcript stability was associated with transient *versus* sustained mRNA accumulation. Together, these findings show that antibacterial responses are organized across multiple regulatory layers and highlight the value of combining nascent and steady-state RNA profiling to resolve the temporal organization of mosquito gene expression.

### *In vivo* transcriptional landscape in *Aedes aegypti* midgut

Although PRO-seq yields highly informative data, its use has been largely restricted to cultured cells, and applications in intact tissues remains limited. The few but notable examples include application in glioblastoma, stool samples, keratinocytes, and *Drosophila* embryos^66–69^. Extending nascent transcription profiling to *in vivo* systems is critical for understanding gene regulation in physiologically relevant contexts. Here, *in vivo* profiling of mosquito midguts revealed tissue-specific transcription and TSN usage, underscoring the importance of studying gene regulation within native context. Expanding such analyses across infection conditions, mosquito strains, and wild populations may provide broader insight into regulatory mechanisms underlying vector competence, while also enhancing characterization of regulatory variation, such as natural promoter variants linked to insecticide resistance and virus susceptibility^70,71^.

### Simultaneous profiling of host and microbiome transcription *in vivo*

Applying PRO-seq to midgut simultaneously captured transcription of host cells and the microbial community at their natural interface, representing, to our knowledge, the first demonstration of host-microbiome co-profiling at the level of nascent transcription. Mosquito midguts harbor a dynamic microbiome that influences vector competence through mechanisms including immune priming, redox regulation, and direct competition with pathogens^52,55^. Accordingly, symbionts such as *Wolbachia* bacteria or insect-specific viruses have attracted increasing interest as tools to interfere with arbovirus transmission^72,73^. In this context, simultaneous measurement of host and microbial transcriptional activity provides opportunities to investigate how microbiome composition and activity shape mosquito transcriptional responses relevant for vector competence.

## Conclusions

In this study, we characterized transcriptional regulation and functional genomics in *Ae. aegypti*, a major vector of arboviruses. Identification of transcription start nucleotides refined genome annotation and revealed the positioning of core promoter elements. Promoters displayed predominantly unidirectional transcription and widespread promoter-proximal Pol II pausing. In addition, we identified transcription at putative enhancers, including highly active enhancer clusters, expanding the regulatory annotation of this important mosquito species.

Antibacterial responses in *Ae. aegypti* were organized into temporally distinct transcriptional waves associated with different biological functions and transcription factors. Rapidly induced waves were enriched for immune genes and antimicrobial effectors, whereas stress-related genes were induced only later. Integration of nascent and steady-state RNA profiling further revealed that gene length and stability contribute to the timing of gene expression programs. Extension of transcriptional profiling to mosquito midguts showed tissue-specific transcription and start nucleotide usage, emphasizing the importance of studying mosquito gene regulation in physiological contexts, and enabled simultaneous profiling of host and microbiome activity *in vivo*. Together, these findings provide new insight into transcriptional regulation, immunity, and host-microbe interactions in *Ae. aegypti*, while generating genomic resources for future studies of vector competence and arbovirus transmission.

## Materials and methods

### Cell culture

The C3PC12 clone of *Ae. aegypti* Aag2 cells^74^ was cultured at 28°C in Leibovitz’s L-15 medium (Invitrogen) supplemented with filter-sterilized and heat inactivated 10% fetal calf serum (Gibco), 50 U/ml Penicilin (Invitrogen), 50 µg/ml Streptomycin (Invitrogen), 1x non-essential amino acids (Invitrogen), and 2% tryptose phosphate broth (Sigma). K562 cells were cultured at 37°C, 5% CO2 in RPMI 1640 + Glutamax (Gibco) supplemented with filter-sterilized and heat inactivated 10% fetal calf serum (Gibco), 50 U/ml Penicilin (Invitrogen), and 50 µg/ml Streptomycin (Invitrogen).

### Mosquito rearing and dissection

*Aedes aegypti* mosquitoes (Black Eye Liverpool strain, BEI resources) were reared at 28°C and 70% humidity with a 12h:12h light/dark cycle. Larvae were fed Tetramin Baby fish food (Tetra) and adult mosquitoes were fed 10% sucrose solution. For midgut dissections, five- to seven-day-old female mosquitoes were dissected as described previously^75^.

### Preparation of heat-killed *E.coli* and stimulation of Aag2 cells

*E. coli* (XL10-Gold) cultures were grown overnight in LB medium and harvested by centrifugation at 6,800 g for 3 minutes. Bacterial pellets were washed twice and resuspended in PBS at OD600 = 1. For heat inactivation, bacteria were incubated at 95 °C for 15 minutes. Heat-killed bacteria were aliquoted and long-term stored at −80 °C. To verify heat inactivation, a sample of the bacterial suspension was inoculated onto LB agar plates and incubated at 37 °C overnight. Efficient heat kill resulted in the complete absence of bacterial colonies on LB agar plates. To stimulate Aag2 cells, heat inactivated *E.coli* were equilibrated to room temperature and the bacterial suspension was directly added to the Aag2 cell culture medium at a final concentration of OD600 = 0.05. Treatment of cells with an equal volume of PBS served as negative control.

### Optimizing PRO-seq for *Ae. aegypti* mosquito cells

PRO-seq libraries were generated from chromatin preparations following previously described protocols^26,27,66^. To optimize library generation from Aag2 cells and reduce adapter contamination, several modifications were introduced as highlighted in Fig. S1A, and the complete procedure is described below.

### Chromatin isolation

Chromatin samples for PRO-seq were prepared according to Himanen *et al.* (2025)^27^, with small modifications. Approximately 30 million Aag2 cells were washed with ice-cold PBS before harvesting with a cell scraper. Cells were pelleted by centrifuging at 400 x g for 5 minutes at 4 °C. After a second wash with ice-cold PBS, cells were resuspended in 1 ml ice-cold NUN buffer (20 mM HEPES pH 7.5, 300 mM NaCl, 1M Urea, 1% NP-40 Substitute, 7.5 mM MgCl2, 0.2 mM EDTA, 1 mM DTT, 20 units per ml RNase inhibitor, 1x Protease Inhibitor Cocktail) and vortexed vigorously for one minute. Subsequently, the chromatin was pelleted by centrifuging at 12,500 x g for 30 minutes at 4 °C. Chromatin was resuspended in 50 mM Tris-HCl, pH 7.5, vortexed briefly and spun down at 10,000 x g, for 5 min at 4 °C. After another Tris-HCl wash, the chromatin was resuspended in chromatin storage buffer (50 mM Tris-HCl pH 8.0, 25% glycerol, 5 mM MgCH3COO)2, 0.1 mM EDTA, 5 mM DTT, 40 units per ml RNase inhibitor) and incubated for 5 minutes on ice. Chromatin samples were fragmented by sonication (Bioruptor pico, Diagenode) with 10 cycles of 30 seconds on / 30 sec off intervals at high intensity settings, flash-frozen in liquid nitrogen and stored at −80 °C. For optimization of PRO-seq for Aag2 cells, chromatin was sonicated for another 5 cycles just before the run-on reaction.

### Run-on reaction

The 2x Run-on Master Mix (ROMM: 10 mM Tris-HCl pH 8.0, 5 mM MgCl2, 1 mM DTT, 300 mM KCl, 40μM Biotin-11-CTP (Perkin Elmer), 40μM Biotin-11-UTP (Perkin Elmer), 40 μM ATP (Thermo Fisher), 40μM GTP (Thermo Fisher), 0.8 units/μl RNase inhibitor, 1% Sarkosyl) was prepared and pre-warmed to 28 °C. Meanwhile, chromatin was thawed on ice, and K562 chromatin was added to Aag2 as spike-in, counted to constitute 1% of total chromatin. To start the run-on reaction, an equal volume of ROMM was added to the chromatin and thoroughly mixed by pipetting up and down, followed by incubation at 28°C for 5 minutes.

### Isolation of nascent transcripts

To terminate the run-on reaction, 500 μl TRIzol LS (Thermo Fisher) and 130 μl chloroform (Merck) were added and samples were homogenized by vortexing, incubated on ice for 1 minute and then centrifuged at 18,000 x g for 5 minutes at 4 °C. RNA in the aqueous phase was precipitated with 100% ethanol, using GlycoBlue (Invitrogen) as co-precipitant, and washed with 75% ethanol. RNA pellets were air-dried for 5 minutes, diluted in RNase-free water and base hydrolyzed with NaOH for 5 minutes, after which the reaction was neutralized with 1 M Tris-HCl, pH 6.8. To remove free biotinylated nucleotides, the RNA was passed through P-30 columns (Bio-Rad). Nascent RNAs, with biotinylated 3’-ends, were purified by Steptavidin Dynabeads (Thermo Fisher) and washed with Binding, High Salt and Low Salt wash buffers (Binding buffer: 10mM Tris-Cl pH 7.4, 300 mM NaCl, 0.1% Triton X-100, High Salt wash buffer: 50 mM Tris-Cl pH 7.4, 2M NaCl, 0.5% Triton X-100 and Low Salt wash buffer: 5 mM Tris-HCl pH 7.4, 0.1% Triton X-100). Subsequently, the nascent RNAs were released from the beads through TRIzol/chloroform-based RNA isolation, precipitated with 100% ethanol and washed with 75% ethanol.

### Preparation of PRO-seq libraries

#### 3’ barcoding and pooling of samples

Sequencing adapters with unique barcodes for each sample (Table 1) were ligated to the 3’ end of the purified nascent RNAs. To this end, 1 µl of 25 µM 3’adapters was added to the nascent RNA, followed by denaturation at 65 °C for 1 minute and incubation in ice water for 1 minute. To each sample, 20 U T4 RNA ligase 1 (NEB) was added and incubated at 25 °C overnight. To remove the non-ligated 3’ adapters, nascent RNA samples were purified with Streptavidin Dynabeads as described above. Afterwards, the barcoded samples were pooled per timepoint per replicate.

**Table 1:**
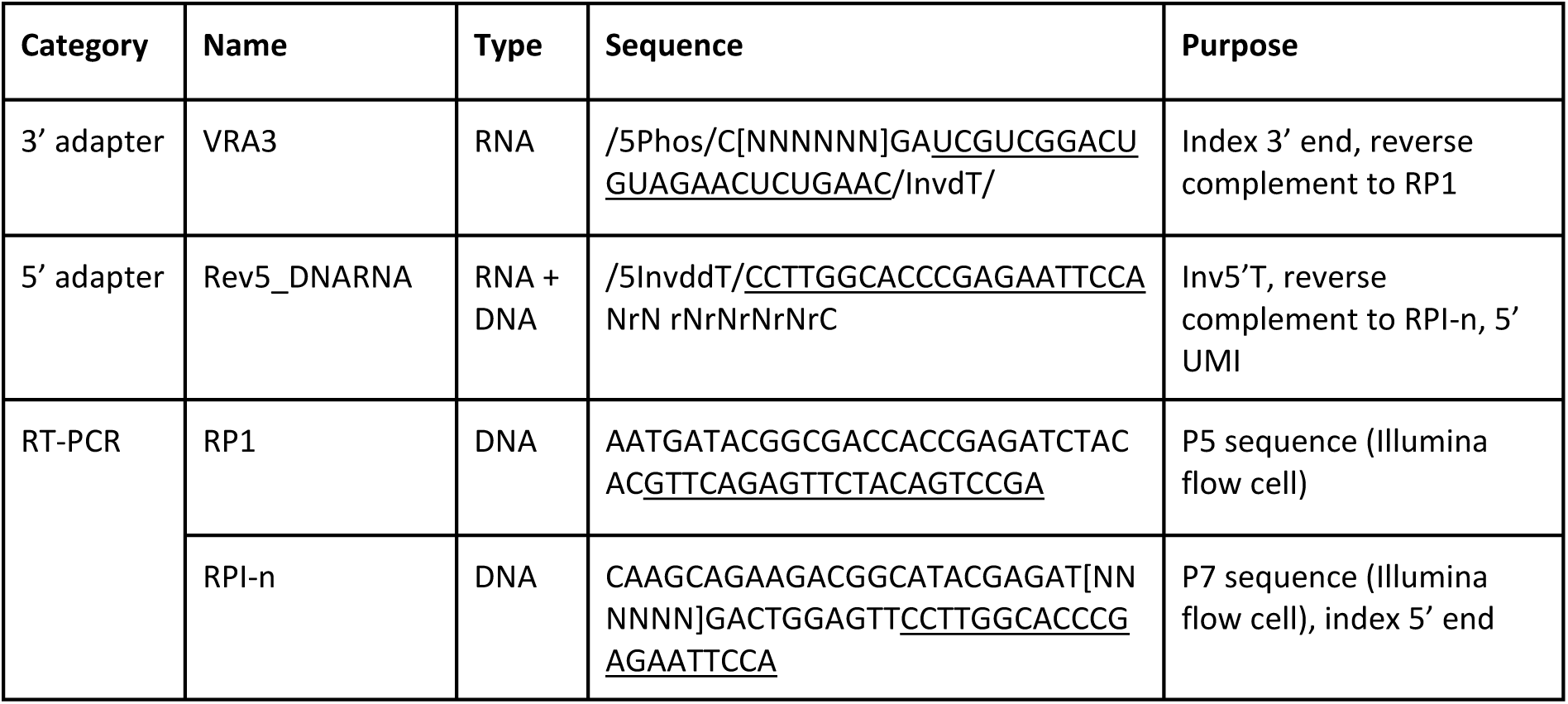
Oligonucleotide sequences.

#### 5’ de-capping, 5’ hydroxyrepair and 5’ adapter ligation

The nascent RNAs bound to Streptavidin Dynabeads were 5’ de-capped by diluting in RNA 5’ Pyrophosphohydrolase (RppH) mix (NEB) and incubation at 37 °C for 45 minutes. Subsequently, 5’ ends were hydroxy-repaired by adding T4 polynucleotide kinase reaction mix (NEB) which was incubated for 45 minutes at 37 °C. For elution of RNA from the beads, the beads were first washed with High Salt and Low Salt wash buffers, and RNA was isolated with TRIzol/chloroform as described above. After air-drying for 5 minutes, the RNA pellets were resuspended in adapter mix with 1 µl of 25 µM 5’ adapter (Table 1) and ATP, and incubated at 65 °C and subsequently in ice-cold water, each for one minute. Subsequently, the ligation mix with 20 U T4 RNA Ligase buffer (NEB) was added and incubated for 3 hours at 25 °C. To remove the non-ligated 5’ adapters, nascent RNA transcripts were purified with Streptavidin Dynabeads and isolated with TRIzol/chloroform, as described above.

#### Reverse-transcription and amplification

The RNA pellets from the previous step, containing the 3’ and 5’ adapter-ligated nascent RNA transcripts, were diluted in a mix containing RP1 primer (Table 1) and dNTPs (Thermo Fisher), followed by denaturation at 65 °C for 1 minute and incubation in ice water for 1 minute. Reverse transcription was performed with Super Script III reverse transcriptase (Thermo Fisher). The reverse-transcribed samples were amplified with Phusion High-Fidelity Polymerase and primers RP1 and RPI-n (Table 1), using the following cycling conditions: 95°C for 2 min; 5 cycles of 95°C for 30 sec, 56°C for 30 sec and 72°C for 30 sec; 5 cycles of 95°C for 30 sec, 65°C for 30 sec and 72°C for 30 sec; final extension at 72°C for 10 min. For library clean-up and size-selection, a 1.4x volume of Mag-Bind® TotalPure NGS beads (Omega bio-tek) was used. Library concentration and size distribution were determined with Qubit dsDNA High Sensitivity Kit and Bioanalyzer High Sensitivity DNA Analysis.

#### Paired-end deep sequencing

The uniquely barcoded libraries were mixed in equimolar volume and sequenced on a NovaSeq6000 to generate paired-end 2 x 51 nt reads.

### mRNA-seq library preparation

mRNA-seq libraries were generated from the same samples as the PRO-seq experiments, corresponding to two biological replicates of *Ae. aegypti* Aag2 cells, stimulated with either heat-inactivated *E. coli* or PBS, for either one or four hours. Libraries were prepared using the TruSeq Stranded mRNA Library Prep Kit (Illumina) and indexed with the IDT for Illumina TruSeq UDI Plate according to the manufacturer’s instructions. A total of 350 ng RNA was used as input per library. Final library concentration and fragment size distribution were assessed using the Qubit dsDNA High Sensitivity Assay Kit (Thermo Fisher Scientific) and the Bioanalyzer High Sensitivity DNA Kit (Agilent Technologies). Libraries were sequenced on an Illumina NextSeq 2000 platform to generate paired-end 2 × 59 nt reads.

### qPRO-seq for midgut tissue

Dissected midguts were homogenized using sterile plastic pestles in 1.5 ml microcentrifuge tubes, each containing 15-20 midguts in 20 µl cold Leibovitz’s L-15 medium. Homogenates were pooled to obtain a total of 50-55 midguts per sample and centrifuged at 10,000 x g for 5 min at 4°C. Pellets were resuspended in ice-cold NUN buffer, after which chromatin extraction was performed as described above. Midgut libraries were prepared following the qPRO-seq protocol (https://dx.doi.org/10.17504/protocols.io.57dg9i6)^53^, using 1 µl of 5 µM 5′ and 3′ adapters for ligation. Final libraries were pooled and size-selected on a 0.5X TBE, 6% PAGE gel stained with ethidium bromide, and fragments from 150 to 650 bp were excised. Gel slices were incubated overnight at 28 °C and 600 rpm in 0.3 M NaCl, 0.1% Tween-20 in Milli-Q. After centrifugation at 5,000 x g for 1 min, the soaking buffer was collected. Residual gel fragments were removed by passing the sample through filtration columns from the GenElute™ Mammalian Total RNA Miniprep Kit (Sigma-Aldrich) to recover remaining eluate. Eluates were pooled and libraries were purified by ethanol precipitation.

### Bioinformatics analysis

#### PRO-seq: de-multiplexing and read trimming

Replicates were demultiplexed based on 5’ barcode by the sequencing facility of the National Genomics Infrastructure (Stockholm, Sweden), after which the R1 of individual samples were de-multiplexed based on 3’ barcode with fastx_barcode_splitter. Corresponding R2 were paired back with fastq_pair. Reads were trimmed for adapters and UMI with fastp. The 3’ barcode was trimmed at the start of R1 with fastx_trimmer and in the case of readthrough, at the end of R2 with cutadapt. The reverse complement of R1 was obtained with fastx_reverse_complement.

#### PRO-seq: alignment and deduplication

Trimmed reads were first aligned to ribosomal genes and their flanking 50 basepairs, after which the mapped reads were discarded from further analysis. The rRNA gene annotation was based on AaegL5.3 gff file version 3. The ribosomal-depleted reads were mapped to the total *Ae. aegypti* genome (AaegL5.3) and the spike-in human genome (T2T-chm13v2). All alignments were performed with bowtie2 --dovetail and only reads in concordant pairs, including multimappers, were kept for downstream analysis. PCR-duplicated reads were removed with UMItools.

#### PRO-seq: file conversions

Mapped and deduplicated reads in bam format were converted to bed files describing the inserts from R1 and R2 together as a single fragment. From these bed files the 5’ and 3’ nt positions were extracted and used to generate bedGraph files with bedtools genomecov. Signal tracks were normalized in two steps: spike-in reads mapping to the human genome were used to correct for sample-specific differences, after which all samples were scaled to the sequencing depth of the PBS control sample of the corresponding experiment.

#### mRNA-seq: alignment and quantification

For mRNA-seq, paired-end sequencing reads were aligned to the *Ae. aegypti* genome (AaegL5.3) using STAR^76^. Exon-levels counts were generated with FeatureCounts^77^ using the annotation from AaegL5.3 gff version 3. Reads were quantified in a strand-specific manner and multimapping reads were retained and fractionally assigned during quantification.

#### Identification of the Transcription Start Nucleotide

Currently annotated transcription start sites were retrieved from AaegL5.3 gff version 3, and regions from TSS - 100 nt to TSS + 400 nt were extracted. PRO-seq 5’ nt reads were pooled per replicate, after which the nucleotide position with the highest 5’ nt PRO-seq coverage was determined (maxPos) for either replicate. Per replicate, only the upper median of max positions were kept to generate a list of Transcription Start Nucleotides (TSN). Transcripts for which the called nucleotide was the same for both replicates, were annotated as the exact TSN.

#### Core promoter elements and sequence logo

For the transcripts for which we defined the TSN with PRO-seq 5’ nt coverage, we generated bed files of 100 nt windows around the currently annotated TSS and around the experimentally identified TSN. Fasta files, generated through bedtools getfasta, were then searched for sense match to the *Drosophila melanogaster* consensus motifs; TATAWAW for TATA box, TAKTY for Initiator and RGWYV for Downstream Promoter Element. To visualize the distribution of their positioning, we used ggplot2 geom_density. This function computes and draws the kernel density estimate as a smoothened version of a histogram^78^. To generate sequence logos, the same 100 nt window bed files were used as input for Weblogo (weblogo.berkeley.edu), which generates and then visualises the position weight matrix^79^.

#### Quantification of Pol II densities at genomic loci

To count PRO-seq 3’ nt reads over genome features, genes were partitioned as follows: divergent (TSS-750 to TSS-100), promoter-proximal (TSS-100 to TSS+100), +1 to +5 nucleosomes (TSS+100 to TSS+1000), remaining gene body (TSS+1000 to CPS-100), cleavage and polyA signal (CPS-100 to CPS+100) and termination window (CPS+100 to CPS+5 kb). The CPS location was retrieved as the transcript end site from the AaegL5.3 file version 3. The TSS locations were updated to TSN according to PRO-seq 5’ nt coverage where applicable. To define transposable elements, we extracted the AaegL5 repeatfeatures from VectorBase and filtered out the following regions: “dust”, “trf”, “simple_repeat”, “rRNA”, “tRNA”, “low_complexity”, leaving all types of transposable elements. Reads within each region were counted with bedtools intersect (see also Figure S3A), using the -s setting to count in a strand-specific manner and -S to count for the opposite strand.

#### Transcription directionality

Transcription directionality metrics were calculated as described by Core *et al*. (2012)^4^. For each TSN, the 100 bp window with the highest PRO-seq 3′ end coverage within ±500 bp of the TSN was identified separately for the sense and antisense strands. Directionality scores (Fig. 2F and 3D) were then calculated as the ratio of sense to antisense reads within these windows. Orientation indices (Fig. S4C) were calculated as the fraction of PRO-seq coverage in the dominant transcriptional direction, ranging from 0.5 (equal sense and antisense transcription) to 1.0 (fully unidirectional transcription).

#### Identification of enhancers and enhancer clusters

Putative transcriptional regulatory elements (TREs) were identified *de novo* using dREG^44,80^ with total PRO-seq 3′-nt coverage as input. While dREG output regions are defined by their start, end, and peak center coordinates, we first standardized all TRE regions to ±500 nt from the peak center before excluding overlap with annotated gene promoters (TSS ±500 nt). Remaining TREs were only retained as putative enhancers if they overlapped with an ATAC-seq and/or H3K27ac ChIP-seq peak called for all three biological samples from the published Aag2 dataset, where peak calling was performed with MACS2 with standard settings and a q value of 0.05, and against an input sample for the ChIP-seq^81^. For each putative enhancer, transcription start nucleotides (TSNs) were defined per strand as the position with the highest 5′ nt PRO-seq coverage within the original dREG interval. Only TSNs concordant between biological replicates were retained. For enhancers with active initiation on both strands, the dominant TSN and strand were determined by the TSN with the highest 5’ nt coverage. Directionality scores and orientation indices were then computed as described above for gene promoters. To enable direct comparison between enhancers and promoters, a control set of gene promoters was generated using the same filtering criteria for gene TSNs (i.e., requiring overlap with a dREG call, overlap with ATAC and/or H3K27ac peak, and replicate-concordant TSN calls). Enhancer clusters were defined following the eClusterer framework (https://github.com/Vihervaara/eClusterer)^27^. Briefly, for those enhancers with defined TSN and eRNA expression ≥5 RPK (reads per kilobase = total enhancer counts / enhancer length in kb), enhancer coordinates were sorted by genomic position, and clusters were formed when ≥5 enhancers resided within a 12.5 kb window. Initial clusters were iteratively extended if additional enhancers were located within 2 kb of the cluster boundary.

#### Visualization of genomic data

Coverage track files were visualized with IGV genome browser^82^. ChIP-seq, mRNA-seq and ATAC-seq bigwig files as obtained from Qu *et al.* (2023)^81^ were RPM-normalized, and PRO-seq 5’ and 3’ nt bedgraph files were spike-in normalized. Metaplots were generated with deepTools computematrix^83^ with bin size settings indicated in figure legends of the corresponding figures.

#### Differential expression analysis

Differential expression analysis for PRO-seq was performed using DESeq2^84^ on spike-in normalized PRO-seq 3′ end reads mapping to gene bodies, defined as “TSS +100 bp’ to “CPS −100 bp’. For mRNA-seq, DESeq2 was applied to exon-level FeatureCounts as described above. For both datasets, differentially expressed genes were identified using p-value < 0.05 and an absolute fold-change > 1.5.

#### Clustering, functional annotation and motif analysis

To cluster and visualize the expression of transcripts upregulated at either 1 or 4 hours post stimulation, k-means clustering was performed and results were visualized as heatmap with R package ComplexHeatmap^85^. Functional annotation analysis was performed for the gene sets associated with each cluster using DAVID Functional Annotation Clustering^86^. Motif enrichment analysis of promoter regions was performed using HOMER (findMotifs.pl)^87^. Promoters were defined as the region from TSN −500 bp to the TSN. For genes with multiple transcript isoforms, the isoform corresponding to the TSN with the highest PRO-seq signal in *E. coli*-stimulated samples was used.

#### Transcript stability

Transcript stability was estimated following the approach described by^48^, by calculating the ratio between steady-state mRNA levels and nascent RNA abundance (mRNA-seq/PRO-seq). To minimize biases associated with long introns and variable elongation rates^48^, both PRO-seq and mRNA-seq reads were quantified over the same genomic space by restricting analysis to exonic regions only. To exclude contributions from promoter-proximal paused Pol II, only reads located >100 nt downstream of transcription start sites were included.

To relate transcript stability to function, protein-coding transcripts were ranked according to their stability score, and the top and bottom quartiles were used for downstream analysis. Functional enrichment of these sets was performed using g:Profiler (https://biit.cs.ut.ee/gprofiler/gost)^88^ for the KEGG pathway database. To obtain stability scores for specific functional groups, genes annotated as zinc finger protein or ribosomal protein (excluding mitochondrial matches) were identified using the ‘description’ field of the AaegL5.3 GFF3 annotation (version 3), and their stability scores were subsequently extracted. Empirical cumulative distribution functions (ECDFs) of stability scores were generated for each group using 1,000 bootstrap replicates. Differences in distributions were evaluated using the Kolmogorov-Smirnov test, and resulting p-values were adjusted for multiple comparisons using the Benjamini-Hochberg procedure.

#### miRNA target prediction

miRNA abundance data for Aag2 cells was retrieved from Miesen *et al.* (2016)^51^. The three most abundant and three least abundant miRNAs were selected for analysis (Table 2). Potential miRNA target sites were identified by scanning transcript 3′ untranslated regions (3′ UTRs) for reverse-complementary matches to miRNA seed sequences. Target sites were classified as 6mer sites (complementary to nucleotides 2-7 of the miRNA), 7mer-m8 sites (complementary to nucleotides 2-8), or 8mer sites (complementary to nucleotides 2-8 with an additional adenosine opposite miRNA position 1)^89^. Transcripts were grouped according to their strongest predicted target site (8mer > 7mer-m8 > 6mer). Empirical cumulative distribution functions (ECDFs) of stability scores were generated for each group using 1,000 bootstrap replicates. Differences in distributions were evaluated using two-sided Kolmogorov-Smirnov tests, and resulting p-values were adjusted for multiple comparisons using the Benjamini-Hochberg procedure.

**Table 2:**
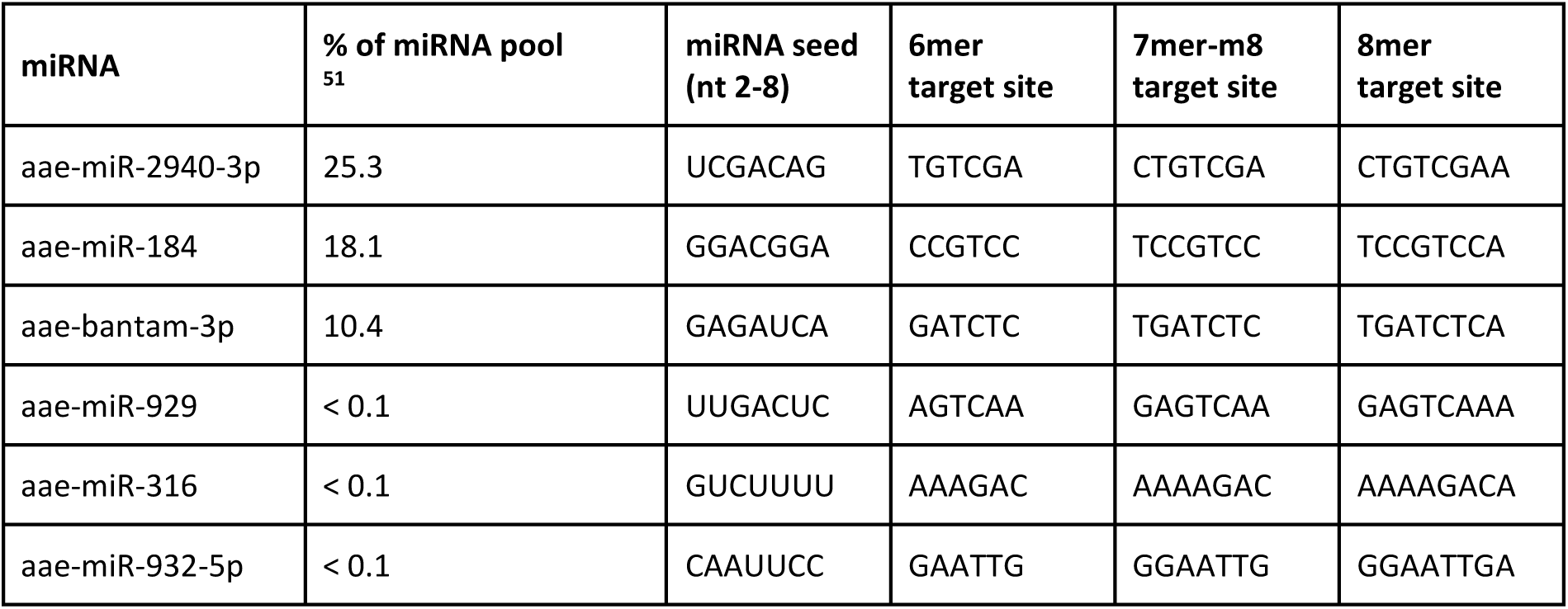
miRNAs and target motifs used for transcript stability analysis.

#### Expression of tissue markers

To assess tissue-specific gene expression, midgut marker genes were retrieved from the *Aedes aegypti* single-cell atlas (https://cells.ucsc.edu/?ds=mosquito)^54^. The top 100 enterocyte markers were selected based on p-value ranking and PRO-seq coverage for these genes was compared between midgut tissue and Aag2 cells.

#### Microbiome analysis

To classify the reads not mapping to the *Ae. aegypti* genome (AaegL5.3), unmapped PRO-seq reads were first filtered to remove sequences shorter than 35 nucleotides. Remaining reads were taxonomically assigned using Kraken2^90^ on the Galaxy server (https://usegalaxy.eu) with the standard Kraken2 database (k2_standard_20210517), after which Bracken^91^ was used to estimate read abundance at the genus level.

To examine PRO-seq coverage over the *Elizabethkingia anophelis* genome, the reads not aligning to the *Ae. aegypti* genome (AaegL5.3) were mapped to the *Elizabethkingia anophelis* reference genome (R26, ASM202366v2) using Bowtie2^92^. Gene annotations were obtained from GenBank (GCA_002023665.2), and operons were predicted with the Operon-.Mapper web server (https://biocomputo.ibt.unam.mx/operon_mapper/)^93^.

## Declarations

## Availability of data and materials

All sequencing data generated in this study have been deposited in the NCBI sequence read archive under BioProject accession number PRJNA1476611 (PRO-seq and mRNA-seq data for Aag2 cells, qPRO-seq data for midguts). The analyzed ChIP-seq and ATAC-seq datasets were obtained from Qu *et al.,* 2023 ^81^ and are available under BioProject accession number PRJNA864851. The mRNA-seq data for dissected midguts is available under BioProject accession number PRJNA933651.

## Authors’ contributions

FAH.v.H., and P.M. conceived the study. FAH.v.H, A.V. and P.M. designed the experiments and analyses. FAH.v.H. conducted the laboratory work. FAH.v.H. and S.H. conducted the computational analyses. All authors interpreted the results. A.V. and P.M. supervised the work. FAH.v.H and P.M. wrote the manuscript, with help from all authors. All authors read and approved the final manuscript.

## Acknowledgements

We thank members of the Miesen and Vihervaara laboratories for valuable advice and discussions. We also thank Milou Stevens, Rebecca Halbach and Ronald P. van Rij for critical reading of the manuscript and Milou Stevens, Jurgen P. Moonen, Oscar M. Lezcano, and Gijs J. Overheul for assistance with mosquito midgut dissections.

