## Supplemental Figures for "Genome-wide nascent transcription profiling uncovers gene regulatory elements and dynamic immune responses in the major arbovirus vector *Aedes aegypti*"

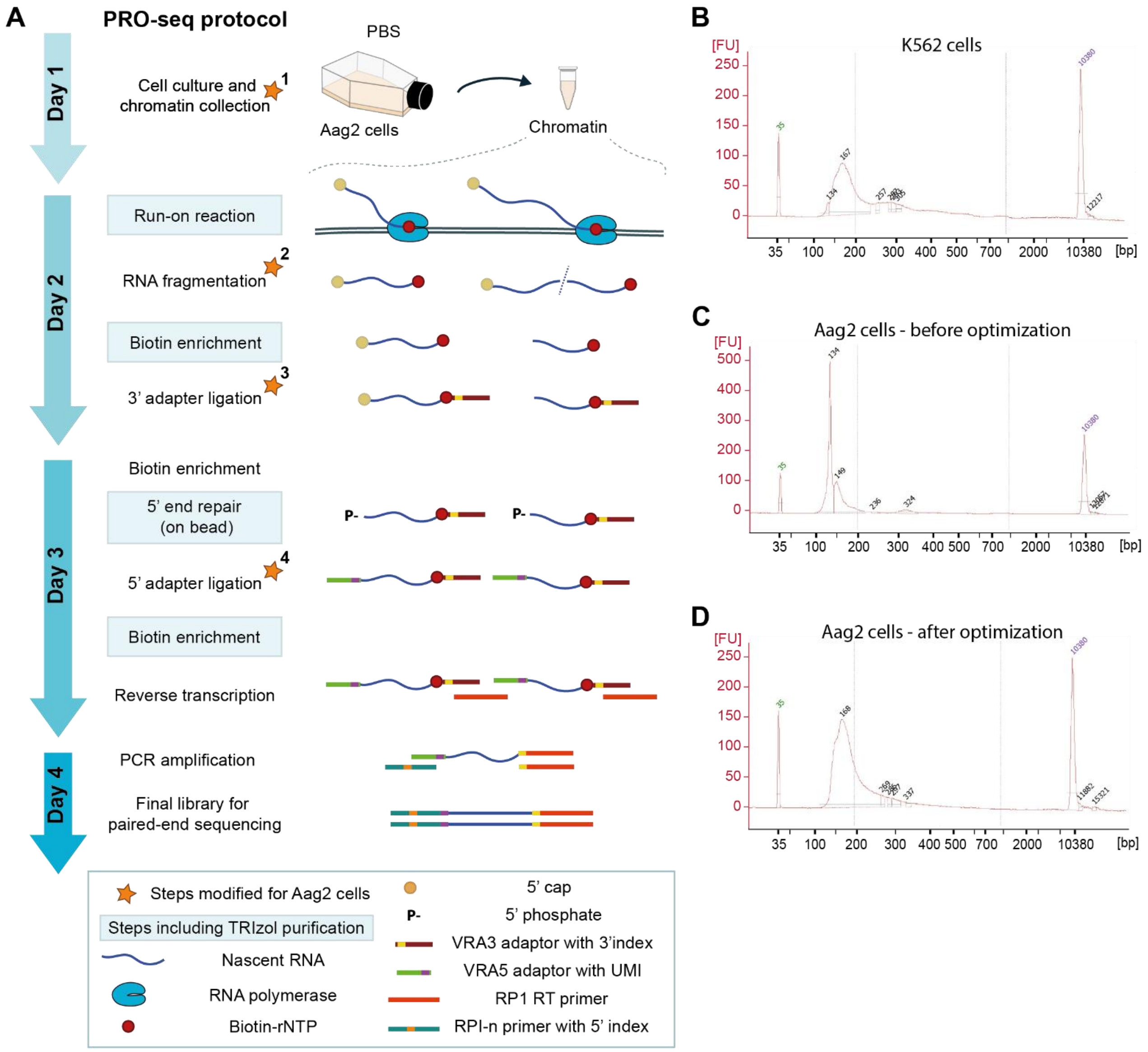


**Figure S1. PRO-seq protocol optimization for *Ae. aegypti* Aag2 cells. A)** Schematic illustration of the PRO-seq protocol. Since its original development by Kwak *et al.* [1], multiple studies have modified the PRO-seq protocol to improve efficiency and throughput. The protocol used in this study was primarily based on the protocol described by Himanen *et al.*[2] with a few additional adaptations for implementation for Aag2 cells. Several of these modifications have also been reported in other PRO-seq workflows[3]. All changes relative to the Himanen *et al.* study are indicated by stars, and the complete procedure is described in the Materials and Methods. The changes include: additional sonication cycles to improve chromatin solubility (*1), omission of the RNA purification step after RNA fragmentation to enhance overall nascent RNA recovery (*2), increased RNA ligase concentration and reduced adapter input (*3, *4), and use of fresh tubes during cleanup steps to minimize adapter carryover. **B)** Size distribution of PRO-seq library generated from human K562 cells as a benchmark. **C+D)** Size distribution of PRO-seq libraries generated for Aag2 cells, before (C) and after (D) optimization of the protocol. Initial Aag2 libraries showed a prominent low-molecular weight peak around 130 nt, indicative of adapter contamination, which was substantially reduced following optimization. DNA fragment profiles in B-D were obtained with Bioanalyzer High Sensitivity DNA Analysis.

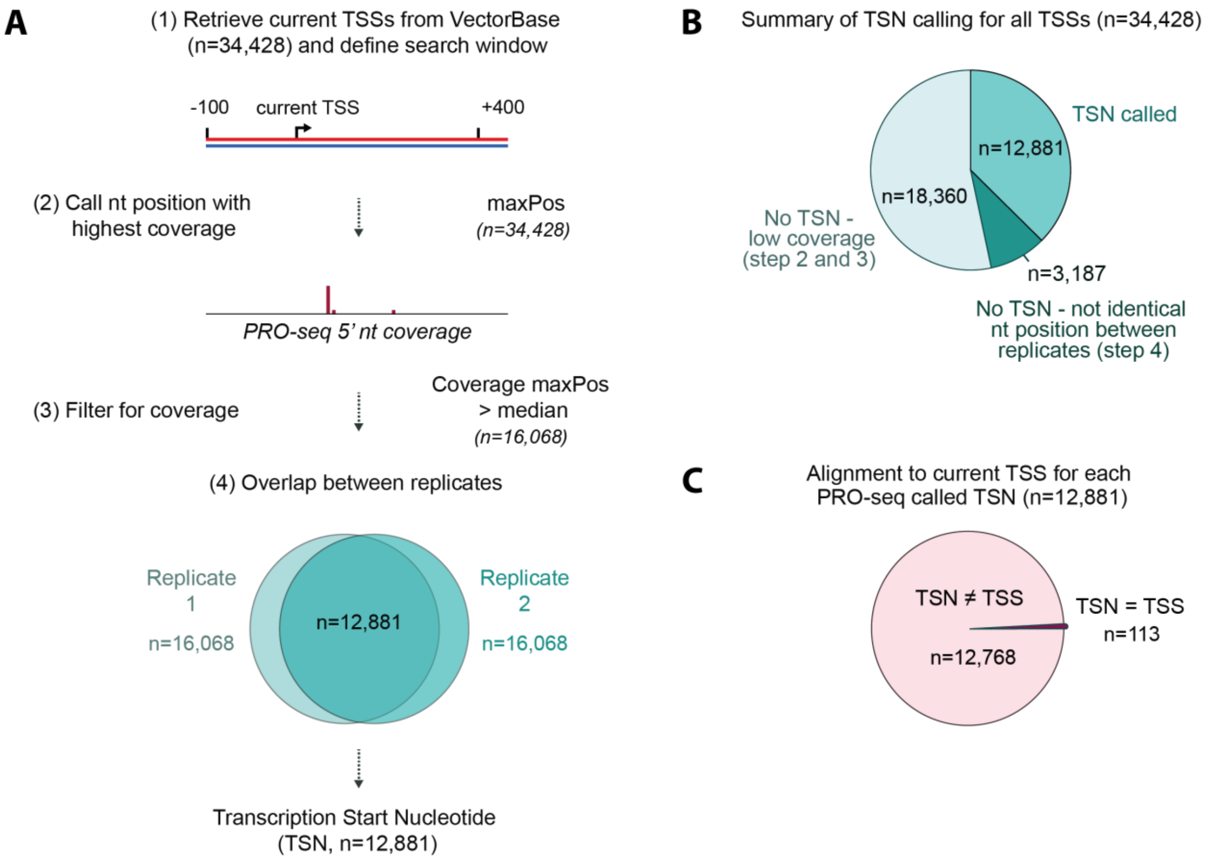


**Figure S2. Strategy for TSN identification at nucleotide resolution based on PRO-seq 5'-nt coverage. A)** Current transcription start sites of *Ae. aegypti* were retrieved from VectorBase, and regions spanning TSS - 100 nt to TSS + 400 nt were extracted (Step 1). Within each region and for each replicate, the nucleotide with the highest 5'-nt PRO-seq coverage (termed maxPos) was identified and regions were ranked based on this coverage; (Step 2). Regions were retained if they were included in the top half of the ranked list in both replicates (coverage maxPos > median; Step 3). Finally, only if the identified maxPos nucleotide position was identical between replicates (Step 4), this was annotated as the exact TSN. All TSN coordinates are listed in Supplementary Table 1. **B)** Summary of TSN calling results based on the procedure described in (A). **C)** Comparison of PRO-seq-defined TSNs to the existing TSS annotation positions to assess alignment at the exact same nucleotide.

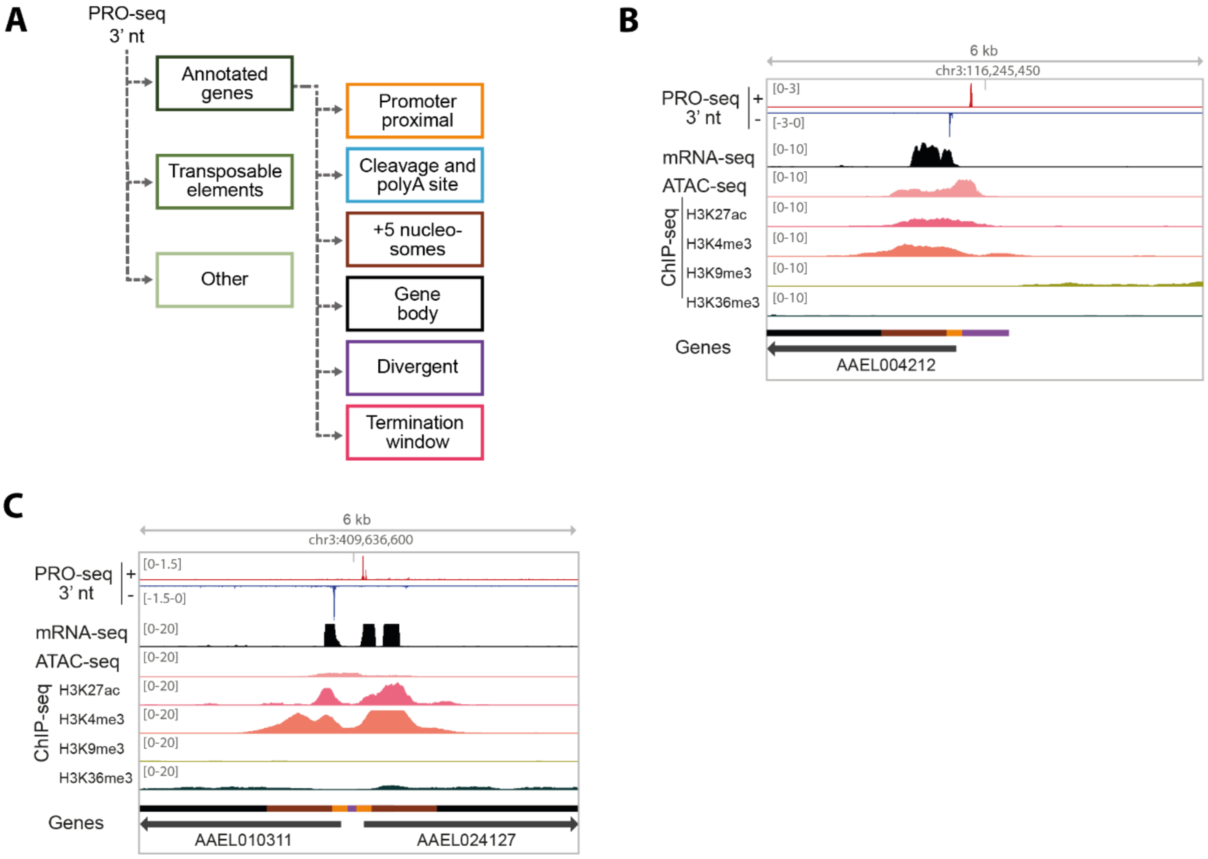


**Figure S3. Genome-wide distribution of transcription across *Ae. aegypti* genomic features. A)** Strategy for hierarchically attributing PRO-seq 3'-nt reads to annotated gene regions (TSN-750 to CPS+5 kb), transposable elements and the remainder of the genome. Reads assigned to annotated genes were further subdivided into functional genomic regions as indicated in Fig. 2A. **B)** Genome browser example of a promoter showing divergent transcription, with PRO-seq 3'-nt coverage for both plus and minus strand. **C)** Genome browser example of a bidirectional promoter, defined by two divergently oriented gene TSNs within 1 kb, with PRO-seq 3'-nt coverage for both plus and minus strand.

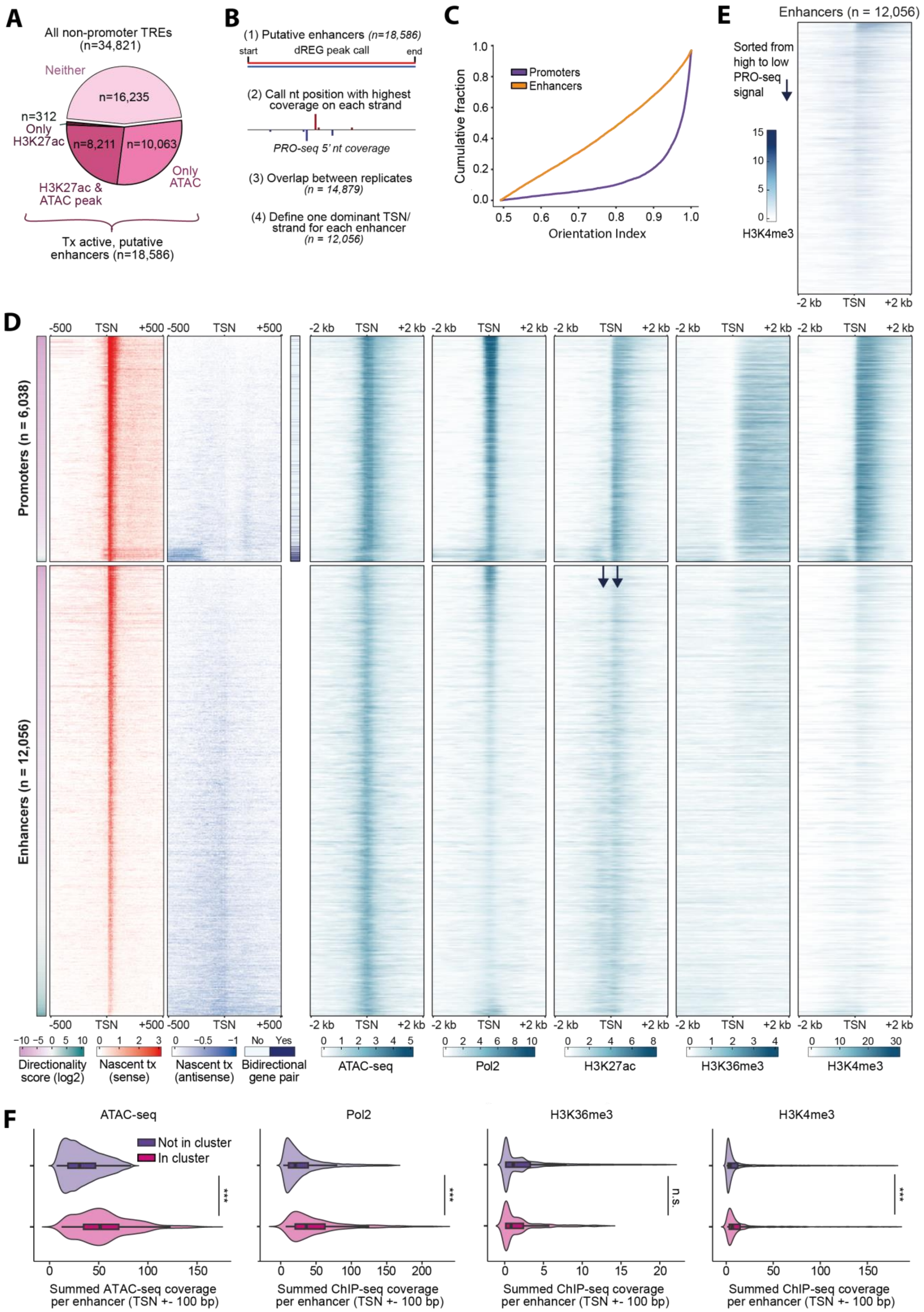
A
All non-promoter TREs (n=34,821)
Neither
n=16,235
n=312
Only H3K27ac
n=8,211
n=10,063
Only ATAC
H3K27ac & ATAC peak
Tx active, putative enhancers (n=18,586)
B
(1) Putative enhancers (n=18,586)
start
dREG peak call
end
(2) Call nt position with highest coverage on each strand
PRO-seq 5' nt coverage
(3) Overlap between replicates (n = 14,879)
(4) Define one dominant TSN/ strand for each enhancer (n = 12,056)
C
Cumulative fraction
Promoters
Enhancers
Orientation Index
E
Enhancers (n = 12,056)
Sorted from high to low PRO-seq signal
H3K4me3
-2 kb
TSN
+2 kb
D
Promoters (n = 6,038)
Enhancers (n = 12,056)
-500
TSN
+500
Directionality score (log2)
Nascent tx (sense)
Nascent tx (antisense)
No
Yes
Bidirectional gene pair
ATAC-seq
Pol2
H3K27ac
H3K36me3
H3K4me3
F
ATAC-seq
Not in cluster
In cluster
***
n.s.
Summed ATAC-seq coverage per enhancer (TSN +- 100 bp)
Summed ChIP-seq coverage per enhancer (TSN +- 100 bp)

**Figure S4. Characterization of putative enhancers. A)** Overlap of non-promoter transcriptional regulatory elements (TREs) with ATAC-seq and/or H3K27ac ChIP-seq peaks, using standardized enhancer intervals (dREG center ±500 bp). **B)** Workflow for defining enhancer TSNs: extract dREG regions (step 1), for each region and each strand call TSN as the nucleotide with the highest 5' nt PRO-seq coverage (step 2), retain only those TSNs concordant between replicates (step 3). As some enhancers contain both a plus and minus strand TSN, assign one dominant TSN (highest coverage) per enhancer (step 4). **C)** Cumulative distribution of promoter and enhancer orientation indices, calculated according to Core *et al.* (2012)[4]: fraction of PRO-seq coverage orientated in the dominant direction, quantifying directionality from bidirectional (0.5) to strongly unidirectional (1.0). **D)** Heatmaps of PRO-seq, ATAC-seq, and ChIP-seq signals (Pol II, H3K27ac, H3K36me3, H3K4me3) at promoters and enhancers (TSN ±500 bp). For all heatmaps the regions are plotted in the same order, according to transcriptional directionality. Note that color scaling differs per mark. **E**) Heatmap of H3K4me3 ChIP-seq signal at enhancers (TSN ±500 bp). Enhancers are ordered from highest to lowest total PRO-seq signal. Note that the color scale differs from panel D. **F**) Violin plots with embedded boxplots; summed coverage of ATAC-seq and ChIP-seq signals (Pol2, H3K36me3, H3K4me3) at enhancers within *versus* outside enhancer clusters, measured as total coverage over TSN +-100 intervals. Values were trimmed to the 5$^{th}$-95th percentile range for visualization; statistical significance was assessed on the full dataset using two-sided Wilcoxon rank-sum tests with FDR correction: ns, FDR ≥ 0.05; *FDR < 0.05; **FDR < 0.01; ***FDR < 0.001.

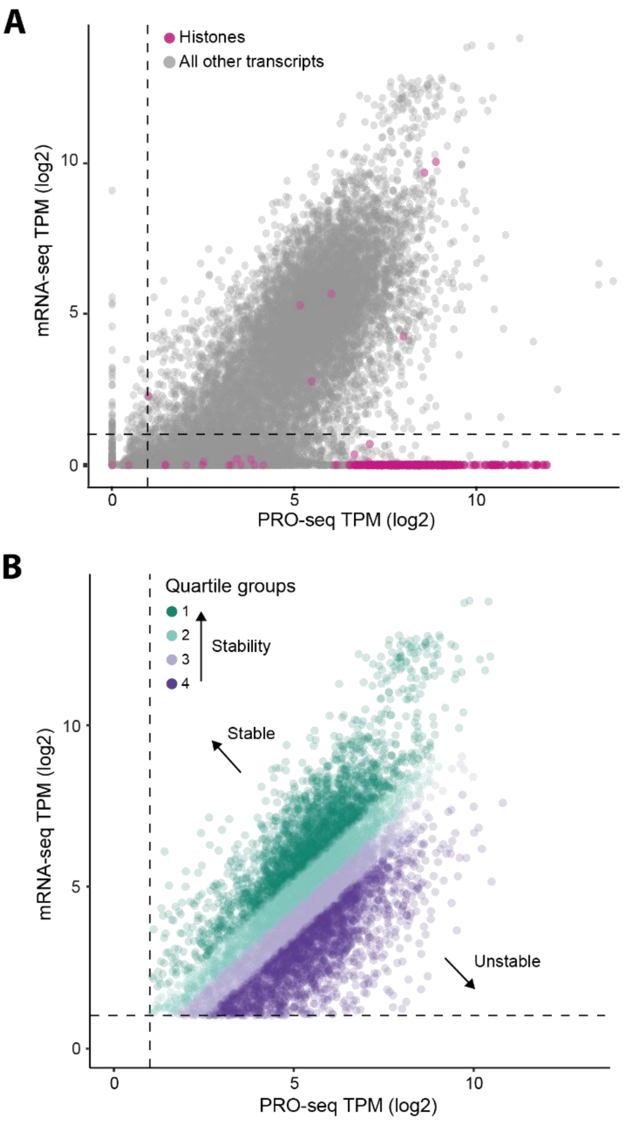


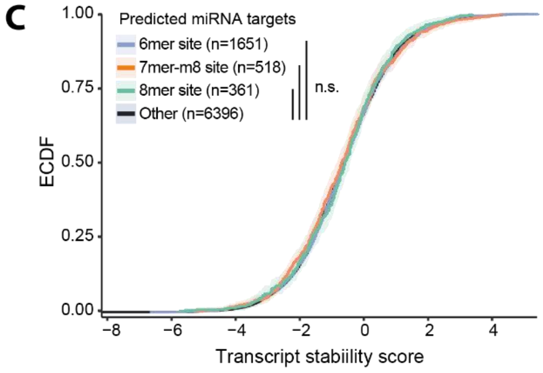


**D** **Stable transcripts (Quartile 1)**

| KEGG | | Stats | |
|---|---|---|---|
| Term name | Term ID | Padj | -log10(Padj) 0–16 |
| Ribosome | KEGG:03010 | $3.122\times10^{-20}$ | |
| Oxidative phosphorylation | KEGG:00190 | $4.695\times10^{-15}$ | |
| Citrate cycle (TCA cycle) | KEGG:00020 | $2.866\times10^{-8}$ | |
| Carbon metabolism | KEGG:01200 | $9.514\times10^{-7}$ | |
| Protein processing in E.R. | KEGG:04141 | $3.170\times10^{-5}$ | |
| Proteasome | KEGG:03050 | $2.950\times10^{-5}$ | |
| 2-Oxocarboxylic acid metabolism | KEGG:01210 | $4.518\times10^{-5}$ | |
| Phagosome | KEGG:04145 | $6.483\times10^{-5}$ | |
| Pyruvate metabolism | KEGG:00620 | $1.193\times10^{-3}$ | |
| Biosynthesis of amino acids | KEGG:01230 | $1.578\times10^{-3}$ | |
| Glycolysis / Gluconeogenesis | KEGG:00010 | $2.117\times10^{-3}$ | |
| Cysteine and methionine metabolism | KEGG:00270 | $5.740\times10^{-3}$ | |
| Nucleocytoplasmic transport | KEGG:03013 | $4.952\times10^{-2}$ | |

**E** **Unstable transcripts (Quartile 4)**

| KEGG | | Stats | |
|---|---|---|---|
| Term name | Term ID | Padj | -log10(Padj) 0–16 |
| Hedgehog signaling pathway - fly | KEGG:04341 | $2.203\times10^{-2}$ | |
| Phosphatidylinositol signaling system | KEGG:04070 | $3.361\times10^{-2}$ | |
| Inositol phosphate metabolism | KEGG:00562 | $4.290\times10^{-2}$ | |

**Figure S5. Transcript stability estimates relate to transcript class, miRNA targeting, and gene function. A)** Scatter plot of mRNA-seq versus PRO-seq in Aag2 cells, shown as log2 transcripts per million (TPM) per transcript (see Methods), with histone transcripts highlighted (violet-red, n = 329). **B)** Scatter plot as in (A), restricted to protein-coding transcripts with log2(TPM) > 1 in both assays. Transcripts were stratified into quartiles according to transcript stability. **C)** Comparison of transcript stability for protein coding genes containing 3' UTR target sites for the three least abundant miRNAs in Aag2 cells. Transcripts were grouped according to target site class (6mer, 7mer-m8, 8mer site) using transcripts without target site as control. **D+E**) KEGG pathway enrichments for transcripts in the highest quartile (most stable, **D**) and lowest quartile (least stable, **E**) of stability scores as depicted in (B).

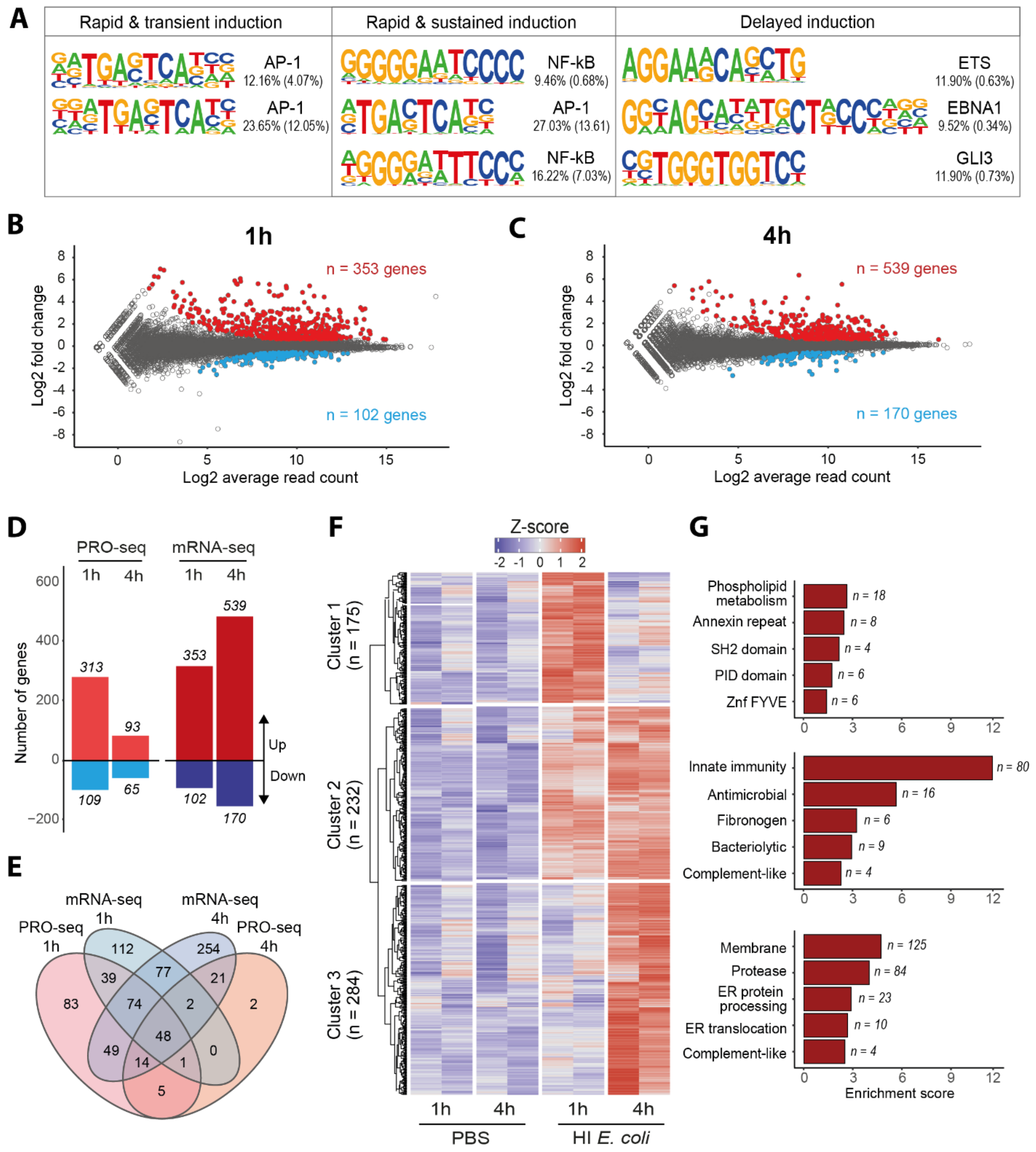


**Figure S6: Regulatory and functional characterization of the Aag2 antimicrobial response. A)** Transcription factor binding motifs enriched in promoter regions of genes from the transcriptional clusters shown in Fig. 5D. HOMER motif enrichment was performed using promoter regions spanning 500 bp upstream of transcription start sites. Motifs with q < 0.05 are shown; total HOMER results provided in Supplementary Tables 10-12. **B+C)** MAplots of differentially expressed genes at 1h (B) and 4h (C) after immune stimulus as measured by mRNA-seq. **D)** Overview of up- and downregulated genes identified by PRO-seq and mRNA-seq. **E)** Overlap of genes upregulated at 1 and 4 hours as identified by PRO-seq and mRNA-seq. **F)** Heatmap showing z-scored mRNA-seq signal of genes with increased mRNA levels at either 1 or 4 hours following stimulation. Genes were clustered using k-means clustering (k=3), with averages of biological duplicates as input. n indicates the number of genes in each cluster. **G)** Functional annotation of the mRNA clusters shown in (F) using DAVID. Top five enriched categories and the top-ranking term per category are shown; total enriched categories and terms are listed in Supplementary Tables 15-17.

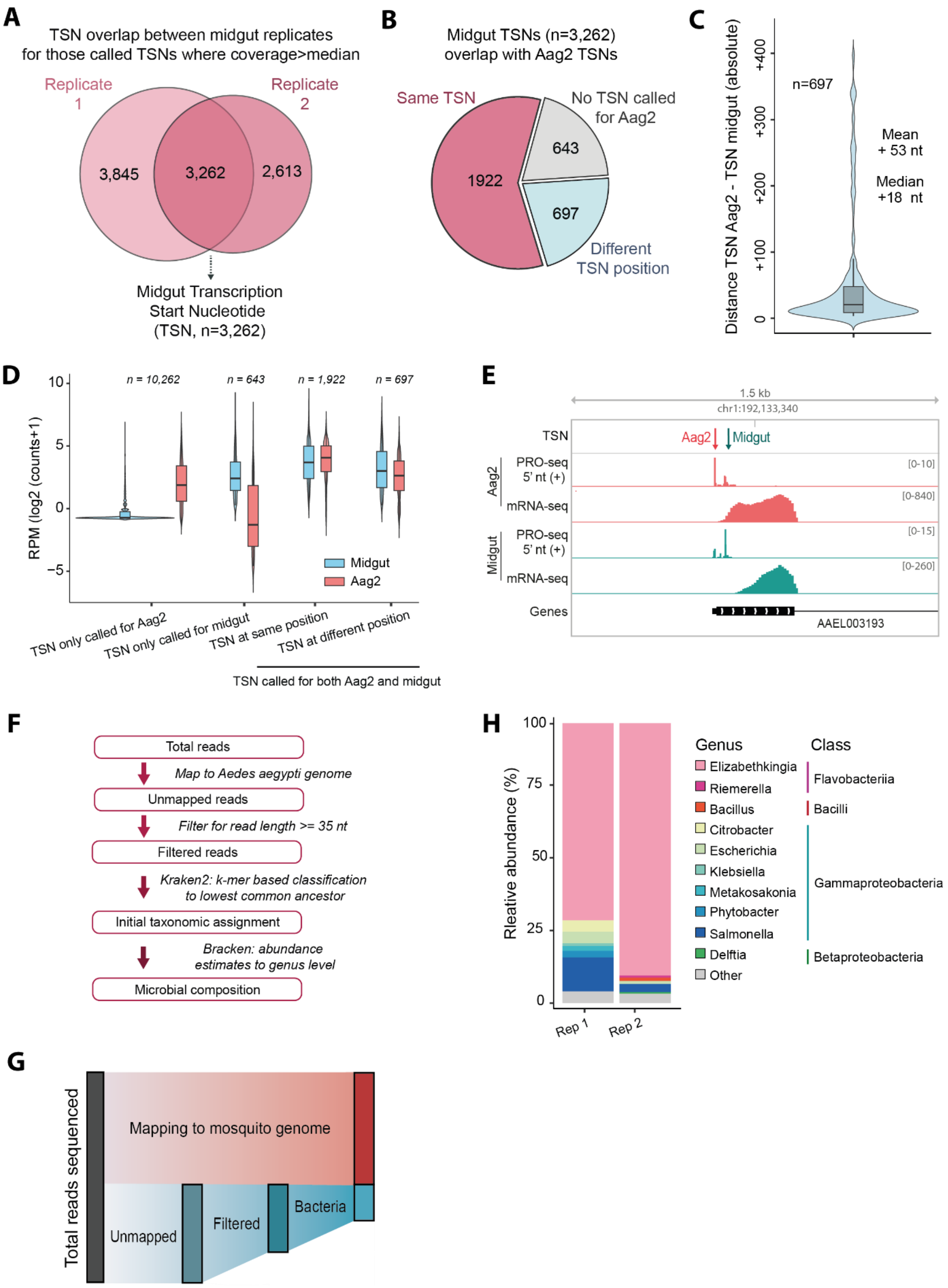


**Figure S7. Tissue-specific transcription initiation and microbiome profiling from midgut PRO-seq. A)** Summary of TSN calling results for midgut PRO-seq data based on the procedure described in Figure S2A. All TSN coordinates are listed in Supplementary Table 18. **B)** Overlap of TSNs called for midgut with TSNs called for Aag2. **C)** Distances between Aag2 and midgut TSNs that differed in position. **D)** Expression levels of transcripts with TSNs defined only in Aag2, only in midgut, or both. The latter group was split into transcripts for which the same or different TSN position was called. **E)** Genome browser view of PRO-seq 5′ nt and mRNA-seq tracks at AAEL003193, showing differential TSN usage in Aag2 versus midgut. **F)** Pipeline for taxonomic classification of reads not mapped to mosquito genome. After filtering out reads shorter than 35nt, remaining reads were classified using Kraken2 and abundance estimated to genus level with Bracken. **G)** Mapping overview of PRO-seq reads from mosquito midguts: reads not mapping to the mosquito genome were length-filtered (≥35 nt) and then taxonomically classified. **H)** Microbiome composition of *Ae. aegypti* midgut as identified by PRO-seq reads (pipeline Figure S7F). Genera with coverage >1000 RPM (0.01%) are shown individually, the rest as "other".
